# Persistent male survival advantage in a protogynous hermaphrodite fish

**DOI:** 10.64898/2026.04.02.716101

**Authors:** Letizia Pessina, Redouan Bshary

## Abstract

In many polygynous species, males face stronger intrasexual competition, higher energetic demands, and lower survival than females, especially under resource limitation or environmental stress. Such sex-specific vulnerabilities are expected to intensify with climate change. Yet, in sequentially hermaphroditic systems, where individuals change sex during their lifetime, how sex and sex change shape survival remains largely unexplored. We studied sex-specific survival and growth in the haremic protogynous cleaner wrasse *Labroides dimidiatus* across eight reefs around Lizard Island, Great Barrier Reef. We tracked a total of 731 adult fish (individually recognizable through marking or idiosyncratic color patterns) over two years. This period included the 2024 El Niño–Southern Oscillation (ENSO), which caused a temporary 1-degree increase in water temperature, severe coral bleaching, and coral mortality at Lizard Island. Contrary to expectations from dioecious systems, terminal-phase males exhibited higher survival than initial-phase females under both normal and in particular ENSO conditions. While male mortality was not affected, female mortality more than doubled during the event, indicating greater physiological or energetic vulnerability. A partial explanation for the overall higher female mortality is their generally faster growth rate, which declined in both sexes during the ENSO event. Our findings challenge existing assumptions of male-biased mortality in polygynous species and highlight that sex and sex change fundamentally shape demographic responses to climate extremes.

## Introduction

In polygynous species, males often experience intense intrasexual competition for access to mates (Casas and Saborido-Rey 2021), resulting in high reproductive skew whereby a small number of dominant males secure most reproductive opportunities (Kruger 2021). This competition imposes distinct life-history trajectories on the sexes (Trivers 1972; Bell 1980): Males typically grow larger to enhance competitive ability, delay maturation to achieve sufficient size, reproduce over fewer seasons, and often exhibit lower survival (Trivers 1972; Parker 1992; Lukas and Clutton-Brock 2014; Hämäläinen et al. 2018; Horne et al. 2020). Elevated male mortality could be partly attributed to their larger body size (Laurie and Brown 1990), which increases energetic demands and vulnerability to resource limitation during periods of food scarcity. These sex-specific costs of reproduction may carry heightened demographic consequences in the context of climate change. Males could prove more vulnerable to the related intensifying and prolonged extreme weather events.

While polygyny is typically studied in dioecious species with fixed sexes, it also exists in protogynous species, where individuals first function as females and only a few larger individuals eventually change sex and become males that then monopolize a harem of females (Ghiselin 1969; Robertson 1974a; Robertson 1974b; Warner and Robertson 1978; Sadovy and Shapiro 1987; Ross 1990; Munday et al. 2006). This type of hermaphroditism is common in coral reef fish, including clades of great ecological and commercial importance, such as parrotfish, wrasses, and groupers (Warner and Robertson 1978; Zhou and Gui 2010; Kuwamura et al. 2020; Sakai 2023). However, how sex and sex change shape vulnerability to environmental extremes remains unknown. A key example of such extremes in the marine environment is the occurrence of the El Niño-Southern Oscillation (ENSO)(Cai et al. 2014; Cai et al. 2021; Cai et al. 2022; I.P.O.C.C. 2023; Thirukanthan et al. 2023; Huang et al. 2024: 20), which causes marine heatwaves and coral bleaching, reducing habitat quality and food availability. Yet, despite evidence of ENSO impacts on corals, fish growth, and mortality (Wellington and Victor 1985; McPhaden et al. 2006; Collins et al. 2010; Ainsworth and Gates 2016; McGowan and Theobald 2017; Hughes et al. 2018; Triki et al. 2018; McClanahan et al. 2019; Triki and Bshary 2019; Huang et al. 2021), its potential for causing sex-specific survival responses in protogynous hermaphrodites remains largely unexplored.

To address this gap, we studied the cleaner wrasse (*Labroides dimidiatus*). This species is a keystone protogynous species essential for maintaining reef fish diversity and abundance through mutualistic cleaning interactions (Bshary 2003; Bshary et al. 2007; Ros et al. 2011; Waldie, Blomberg, Cheney, Goldizen, and Alexandra S. Grutter 2011; Ros et al. 2020). It has also emerged as a model species to study the effects of global change on behavior and cognition (Triki et al. 2018; Paula et al. 2019; Pereira et al. 2025). Both the behavior and survival of this species were previously documented to have been negatively affected by extreme weather events (Wismer et al. 2014; Pereira et al. 2015; Triki et al. 2018; Triki et al. 2019; Pereira et al. 2025), but without studying potential sex related differences.

We tracked individually marked fish across eight reef sites around Lizard Island, Great Barrier Reef, over a two-year period that encompassed the 2024 ENSO event. The 2024 ENSO led to sustained and extreme warming during the austral summer (NOAA 2019; Henley et al. 2024; Australian Institute of Marine Science 2025), resulting in the most severe coral cover decline on record at Lizard Island (Australian Institute of Marine Science 2025). This extreme climatic event provided a natural experiment to examine sex-specific adult survival in a protogynous species not only under ‘normal’ conditions but also under environmental stress. We measured growth and survival, contextualized with in situ coral bleaching surveys as well as published water temperature measurements, to describe general conditions and to confirm the negative effects of ENSO in our research area. Contrary to expectations based on male-biased mortality in other polygynous species, our results reveal that male cleaner wrasse exhibited lower mortality than females. These findings challenge existing assumptions about sex-based vulnerability and highlight the need to account for hermaphroditic systems in assessments of climate sensitivity and population dynamics.

## Methods

### Temperature

To characterize water temperature at Lizard Island, we used lagoon temperature data collected by AIMS sensor buoys deployed at Lizard Island (northern Great Barrier Reef) between July 2022 and November 2024(Australian Institute of Marine Science (AIMS) 2020).

### Site and study populations

The study took place at Lizard Island, on the Great Barrier Reef, Australia, in 2023 and 2024. To determine seasonal differences and detect the effects of El Niño, data were collected in Summer (February to April) and winter (July to September) for both years. Eight sites were selected as our study populations based on their varying cleaner and client densities (Fig S5). Fish were caught while SCUBA diving using hand nets (10 × 15 cm) and barrier nets (4.7 × 1.8 m or 1 × 1.2 m). To facilitate individual identification, we used visual implant elastomer (VIE) tags, a well-established methodology in small reef fishes (Jungwirth et al. 2019). Tagging was conducted underwater, with each fish kept in a hand net to maintain normal saltwater flow. From initial capture to release, the entire process took less than 2 minutes per fish. All individuals resumed normal behavior immediately after release. No non-target cleaner fish were accidentally captured.

Each adult focal individual received two injections, with each injection placed at one of four possible body locations: the front and back of both the left and right sides of the body (Fig S6). Injections were applied to the clear tissue, specifically the white or beige band located above the black band. Each injection used one of six possible colors (red, pink, yellow, green, blue, or white). This tagging scheme, which combines 2 tags, 4 locations, and 6 colors, allows for up to 1,296 unique codes. This number increases further if the order of color placement is considered (e.g., red followed by yellow is distinct from yellow followed by red).

Juvenile focal individuals, typically around 40 mm in length, received only one injection, using colors most visible on their darker body coloration (red, pink, or yellow), resulting in 108 possible unique tags.

In some cases, tagging was not necessary, as individuals could be reliably identified based on unique physical features such as darker or lighter pigmentation, irregularities in the lateral black band, or other distinct markings (Fig S7).

VIE tags are created using a two-part mixture consisting of a viscous pigment and a hardening agent. The hardener helps the tag maintain its structural integrity as the fish grows, preventing the gradual fading of the pigment (Northwest Marine Technology, Inc. 2017). However, because the hardening agent sets rapidly at high temperatures, causing the entire syringe to solidify and go to waste, we chose not to use it. Given the large number of fish that needed to be tagged in this study, this decision minimized material loss. Nevertheless, throughout the study, we observed no loss of marking. While the absence of a hardener occasionally led to tag expansion as the fish grew, all tags remained clearly visible throughout the observation period. To further confirm the durability of our VIE tags, a subset of previously tagged individuals was opportunistically recaptured 12-20 months after marking and examined. All retained a clearly visible tag. These observations were used solely to verify tag durability and are not otherwise analyzed in this manuscript. Overall, our VIE tags demonstrated greater long-term retention than previously reported (Jungwirth et al. 2019)-

Except for two nearby sites (Mermaid West and Mermaid East), all sampling locations were geographically isolated coral reef patches, separated by broad sandy areas or open water. As the cleaner wrasse is a territorial reef fish that does not cross open water, migration between sites is not possible. Therefore, VIE tag combinations could be reused across different sites. However, at the two closely located Mermaid sites, VIE combinations were not repeated to avoid any potential for tag overlap due to possible movement between those locations.

This study is part of an ongoing long-term monitoring project that began in July 2022. The primary researcher participated in every field trip and, to date, has conducted approximately 1,000 dives, totaling around 1,750 hours of direct underwater observation time. This effort involved repeated monitoring of the same individual fish, resulting in an exceptional level of familiarity with their appearance and behavior. This intensive, long-term monitoring enabled individual recognition, including the fish that lacked tags but instead showed peculiar patterns.

Between 2022 and 2024, the dataset included 731 adult fish (120 untagged) and 495 juveniles (291 untagged). Age class was based on life stage at initial observation. Not all individuals were included in every analysis due to logistical constraints during data collection. For example, size measurements were sometimes limited by equipment battery life. Consequently, sample sizes vary across different datasets and analyses.

### Sex identification

Sex identification was based on behavioral observations in the field, a widely accepted method in fish behavioral ecology. In this species, previous research has documented distinct sex-specific behaviors that reliably distinguish males from females (e.g., territorial displays, courtship roles), and these behavioral indicators have been validated through physiological assessments, including gonadal inspections (Robertson 1974b). Thus, behavioral sexing in this study was grounded in well-established and independently verified criteria.

Several qualitative behavioral differences were used here for sex identification : (i) Flutter-Run, a rapid display involving tail fluttering and fin spreading, performed exclusively by males as they swim past females while presenting a lateral view (Robertson 1974b); (ii) Sexual signaling, where females respond positively to male courtship advances. This includes the Body-Sigmoid, a static display in which the female sharply curves her body into an S shape, with the belly bulging toward the male, typically as a signal of readiness to spawn (Robertson 1974b); (iii) Coloration differences, as females develop a distinct sexual color pattern during courtship, whereas males do not (Robertson 1974b); and (iv) Spawning posture, where males adopt a superior position during the later phase of courtship, such as the upward spiral, straddling the female and leading the upward rush (Robertson 1974b).

In addition to these qualitative traits, more quantifiable behavioral differences were also used. Males showed greater tolerance of females in proximity, particularly around their feeding areas, and exhibited a higher rate of movement within their territories, frequently visiting females and patrolling territorial boundaries (Robertson 1974b).

Sex identification was further confirmed through spawning patterns. All individuals classified as males were observed spawning with smaller partners, while all females were seen spawning with larger partners. Spawning invariably involved only two individuals, which excludes the existence of sneaker males that would join a spawning event to add their sperm. Over the first year of data collection, 42 individuals were confirmed to have changed sex. This was determined by observing each individual initially spawning with a larger partner (as a female) and later spawning with a smaller partner (as a male). Previous research has shown that when the dominant male disappears, the sex-changing female immediately begins to exhibit male-typical behaviors to establish dominance and secure her new position (Robertson 1974b). Although the physiological process of sex change may take longer to complete, behavioral cues provide a reliable and rapid means of identifying ongoing sex change events.

### Growth rates

We assessed the growth of each focal fish using a calibrated underwater stereo-photogrammetric system, consisting of two GoPro Hero 8 cameras mounted on a fixed rig (Seager 2006). Footage was processed with EventMeasure (Seager 2006) which enables precise three-dimensional measurements based on synchronized stereo images. Before data collection, the system was calibrated using CAL, a dedicated software that ensures accurate geometric alignment between the camera pair (Seager 2006). This stereo setup significantly improves measurement precision over traditional visual estimations (Michael et al. 2011). In the first three seasons, size measurements were taken every two weeks, whereas in winter 2024, they were collected at a reduced frequency of once a month.

The error of this software is known to be around 1-2mm (Euan et al. 2010). To further assess the accuracy of the stereo camera system for our study, a calibration bar with three known distances was used to measure the system’s error. Measurement error was ± 1.13mm using the calibration bar and ± 1.81mm of wild cleaner fish due to movement and growth between measurements (See supplementary materials 7 for more details).

Measurements occasionally indicated that an individual had shrunk since the last assessment. To address this likely consequence of a measurement error, we adopted a standard approach and retained the previous sizing in such instances, adjusting the growth to 0.

To evaluate the potential impact of adjusting negative growth values, we ran an additional model (Model 3.3; Supplementary materials 8) on our raw growth dataset without correcting for shrinkage. The predicted distribution was nearly identical to the predictions from the primary analysis (Model 3.2), indicating that correcting for negative values did not bias the overall pattern (Fig S9)

### Survival

The presence and absence of focal cleaners were updated biweekly during each field season. Survival was assessed strictly within each field period and does not account for intervals between field seasons. At the beginning of each study period, any unmarked adult individual encountered within the study area was captured and tagged, ensuring full control over the composition of each deme from the outset. When an individual was no longer observed in its territory, nearby harems and surrounding reef areas were checked to confirm disappearance or to continue their sampling. Movements between harems were consistently detected in real-time from the beginning of the study, and individuals who shifted harems were continuously tracked. Only in three rare cases were individuals initially considered missing and later rediscovered. These were younger females that had migrated to previously uninhabited peripheral areas, rather than entering existing harems, which later proved to be successful strategies for initiating sex change. These exceptions did not undermine the general approach, as movements within the monitored network were always promptly identified. The appearance of a new cleaner within a previously occupied territory further supported the conclusion that the original individual had disappeared.

For each focal individual, sex and life stage were recorded and updated at the beginning of each study period.

### Fish census

Cleaner fish density at our study sites was initially determined by conducting 10 replicate 30m transects per site. Subsequent fish counts were then based on a more sophisticated methodology that included all reef fish, except for nocturnal and cryptic species. These counts are used in supplementary materials 2 for an additional analysis of cleaner fish’s client densities. Counts were conducted in the Australian summers and winters of 2023 and 2024 and stratified into the two habitats of the reef in which our focal cleaner resided: The reef crest and the reef base. The reef crest is defined as the seaward edge of the reef flat (Green 1994), and the reef base as the bottom of the reef slope, where it joins the sand flat (Green 1994). Each fish census involved five 30-meter transects, spaced 2 meters apart, running parallel to the reef edge within each habitat for each site. Counts were conducted during three swims: i) the first swim was used to count large fish swimming above the reef within a 5m belt, ii) the second to count medium-sized fish swimming on the reef within a 3m belt, iii) the last one to count fish species of the family Pomacentridae within a belt of 2m. The 3m belt width for medium-sized fish was selected based on a previous study demonstrating its effectiveness in counting wrasses(Green 1994), and it is also the one from which we extracted our cleaner wrasse counts. Each transect swim was preceded by a 2-minute wait to reduce the impact of diver disturbance from the previous count. Each transect was swum at a constant speed and performed in approximately 10 minutes. Only individuals larger/equal to 4cm were considered in our counts. Fish were identified at the species level, following the WORMS nomenclature, whenever possible. A total of 343 species and 51 families were encountered over the years.

### Statistical Models

Data analyses were conducted using R version 4.3.1(RStudio Team 2020). The different research questions were addressed using various statistical models, including Generalised Linear Mixed Effects Models (GLMM), Linear Models (LM), and General Linear Models (GLM). These analyses were performed using the “lme4” (Bates et al. 2015), “nlme” (Pinheiro 2011), “Agricole” (de Mendiburu 2023), and ‘stats’ (R Core Team 2023), packages in R. Diagnostic plots and residual analyses were employed to evaluate the fit of the chosen models. Post-hoc analyses were conducted with Tukey’s adjustment method using the “emmeans” package (Lenth 2023). Full model specifications are available in Supplementary Materials 9. Below, we outline the purpose of each model

Our analyses of cleaner wrasse life history focused on patterns in density, growth, and mortality.

We first examined changes in adult cleaner wrasse density (total length >=5 cm) across years and seasons (Model 1), using standardized fish counts per 100 m² at each site. Density values were log-transformed to normalize the distribution. We modelled these data using an LMER with fixed effects for season, year and their interaction. A random effect for site was included to account for site-specific variability.

To compare growth rates across seasons and years, the cleaner fish population was subdivided into 10 mm size classes to account for ontogenetic differences in growth (younger fish grow faster). The growth rate for each individual was calculated by subtracting the initial size from the final size, then dividing by the number of days between measurements, to obtain a per-day growth rate. Individuals with less than one week between measurements were excluded. Only fish in size classes 50 mm, 60 mm, 70 mm, and 80 mm were included in the analysis due to consistency of sample sizes across seasons.

Growth was analyzed using two complementary modelling approaches. First, we modelled log-transformed growth as a Gaussian response in an LMER (Model 2.1) to assess the effects of size class, season, and year, including their interactions. Random intercepts for site and individual ID were included to account for spatial variability and repeated measures on individuals.

Second, we used a GLMM with a Tweedie distribution (Model 2.2) to model growth rate as a function of sex, year, and season, including their interactions. This model incorporated random intercepts for individual ID and site to address repeated measurements and spatial heterogeneity, and allowed for dispersion to vary by year to accommodate changes in variance over time. The Tweedie distribution was selected to better handle the data’s distributional characteristics, including zero inflation and continuous positive values. Both models focused on data where negative growth values had been adjusted.

Finally, mortality was investigated, focusing on adult males and females, using an LMER (Model 3), with the response variable transformed by the arcsine square root to meet model assumptions. The model included fixed effects for sampling period, sex and their interaction. A random intercept for site was included to account for spatial clustering. To address heteroscedasticity across sampling periods, a variance structure allowing different residual variances per sampling period level was incorporated via the “varIdent” weighting function.

## Results

### 1. Ecological variables

Environmental data confirmed that the austral summer (2023) experienced no particular thermal stress or bleaching (NOAA 2019). On the contrary, the 2024 ENSO resulted in sustained and extreme sea surface temperatures across the Great Barrier Reef (GBR), with Degree Heating Weeks (DHWs) exceeding 8, a threshold associated with a high risk of coral bleaching (NOAA 2019). This heat exposure coincided with the fifth recorded mass bleaching event on the GBR and the fourth global bleaching event (Australian Institute of Marine Science 2025) (Fig S1) At Lizard Island, these conditions corresponded with widespread coral bleaching and substantial mortality of hard corals. This stark difference provided a clear environmental context for evaluating survival and growth of cleaner wrasse males and females under normal conditions and under thermal disturbance. The latter was corroborated by supplementary photo-quadrat surveys showing consistent patterns of bleaching and coral mortality at our study sites (Supplementary Materials 2, Fig S2).

### 2. Cleaner wrasse performance

There was a significant decrease in adult cleaner wrasse density in 2024 compared to 2023 (Type II, Wald Chi-square Test: Chisq = 8.32, df = 1, p = 0.004). There was no significant seasonal difference (TL >= 5 cm, Type II, Wald Chi-square Tests: Chisq = 0.34, df = 1, p = 0.56) and no significant interaction between year and season (Type II, Wald Chi-Square Test: Chisq = 0.25, df = 1, p = 0.62; Fig 1). On average, cleaner wrasse densities declined by 17.44% in 2024 relative to 2023.

**Fig 1.**
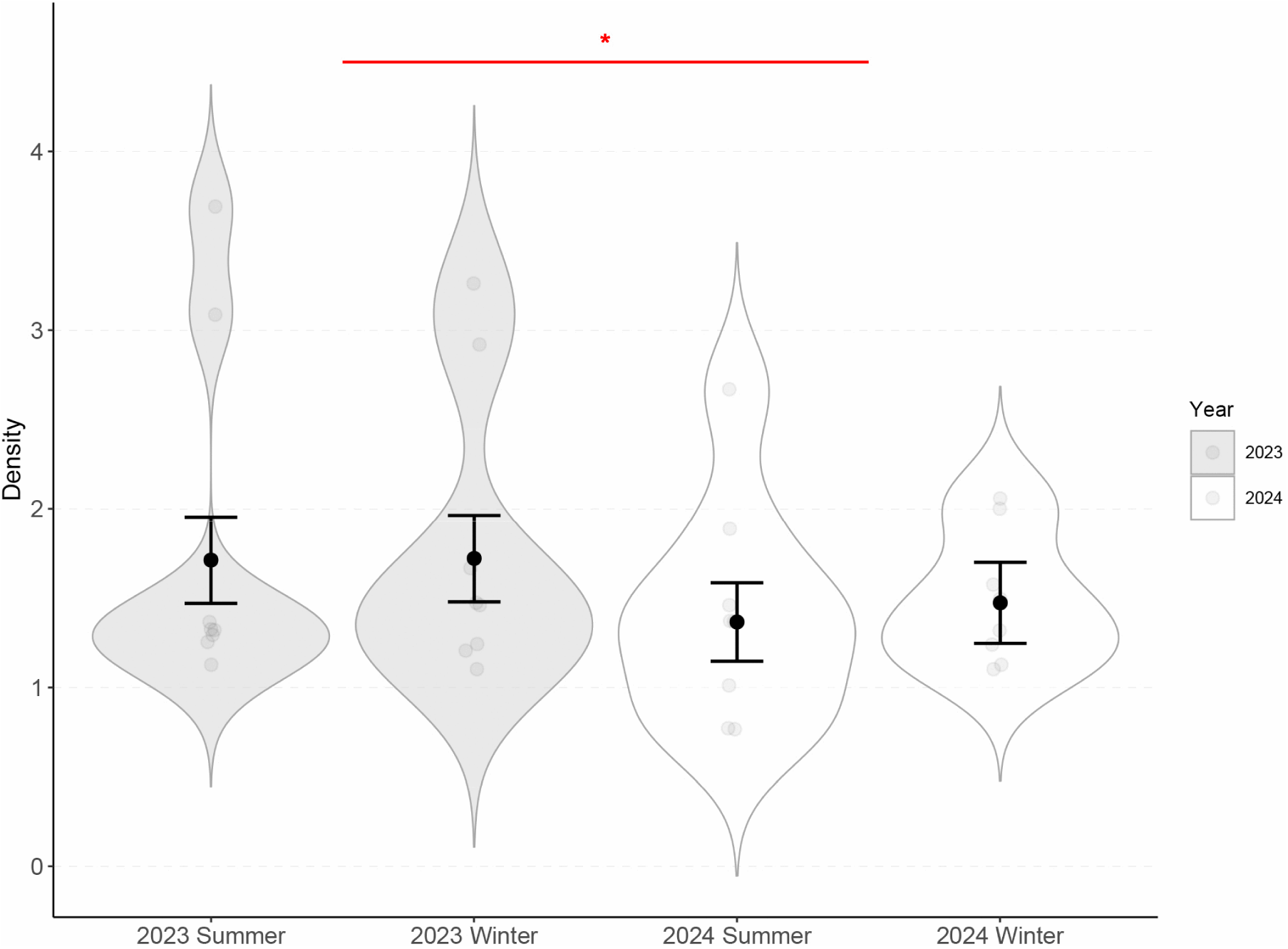
Cleaner fish density before, during, and after the 2024 ENSO event. Distribution of the cleaner wrasse density (individuals per 100 m²) across study periods shown as violins. The jitter represent raw site-level data while overlaid points and error bars represent model-estimated means ± standard errors (SE). Each one of our 8 sites is represented by a single jitter point for each study period.

Two years of monitoring marked individuals have shown that adult cleaner fish are highly site-attached, with only a small proportion relocating to neighboring harems. In the first year alone, only 12.7% of females were observed migrating to adjacent harems. These movements were consistently detected in real time and hence were not mistaken for disappearances. Thus, when an individual was no longer observed, it could reliably be interpreted as a mortality event. With this in mind, our analysis revealed that mortality rates differed between the sexes, both overall and across seasons. Specifically, female cleaner wrasses exhibited an overall higher mortality than males (Fig 2; Type II tests, Wald Chi-square Test: Chisq = 17.33, df = 1, p < 0.0001). During the baseline period (2023 Summer), females experienced an estimated 7–8 times higher mortality rate than males, according to back-transformed values from the mixed-effects model. Mortality also varied significantly across study periods (Type II tests, Wald Chi-square Test: Chisq = 8.1, df = 3, p = 0.04), and there was a tendency for a significant interaction between sex and study period (Type II tests, Wald Chi-square Test: Chisq = 7.58, df = 3, p = 0.06; Fig 2).

**Fig 2.**
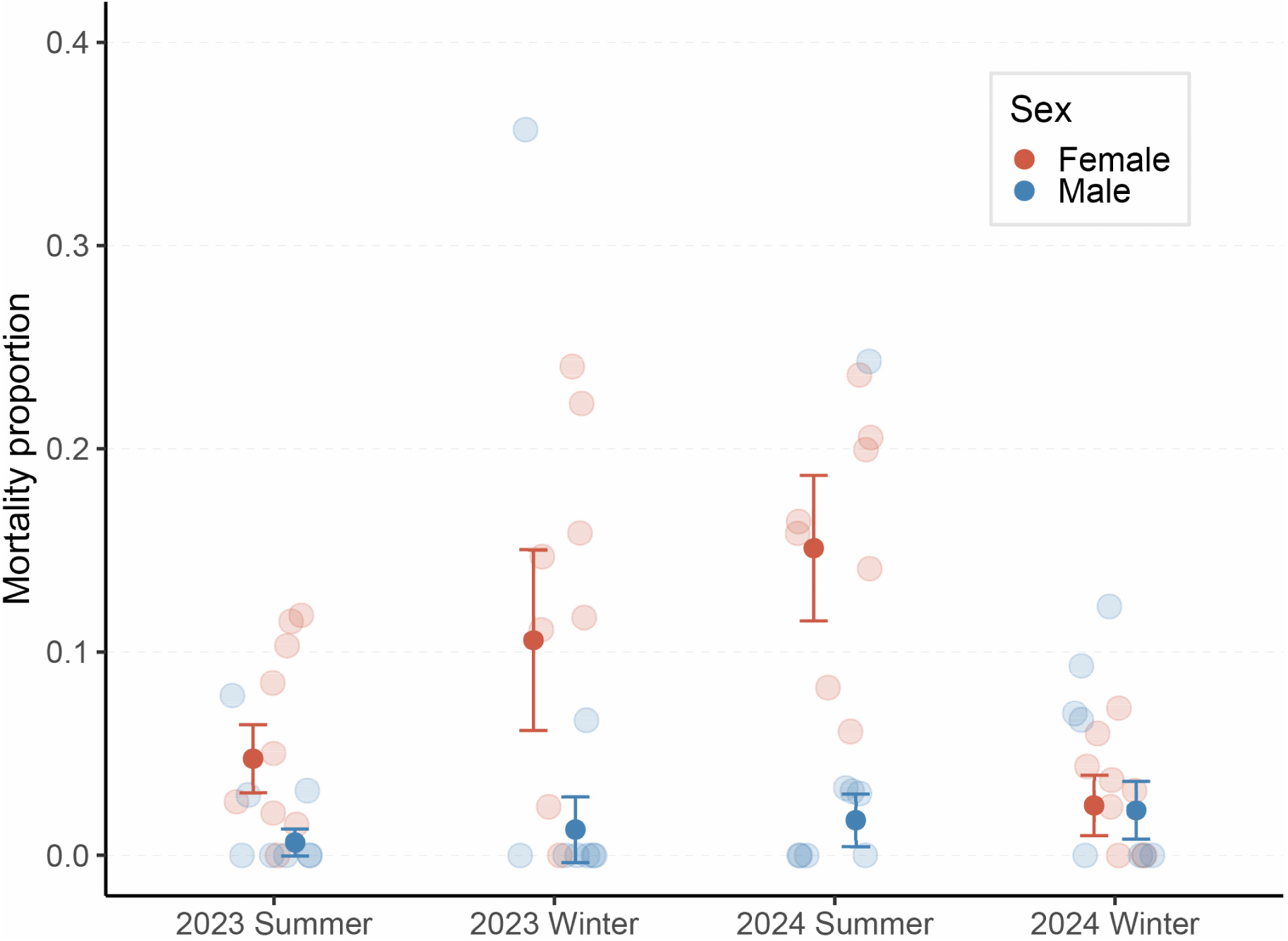
Sex-specific mortality rates of cleaner wrasses before, during, and after the 2024 ENSO event. Jitter point of the monthly mortality rates of male and female cleaner wrasse across three study periods: pre-ENSO (2023), ENSO peak (2024 summer), and post-ENSO (2024 winter). Overlaid points and error bars represent model-estimated means ± standard error (SE). Mortality levels were estimated for each sex at each site.

Post hoc analyses showed that female cleaner wrasses consistently experienced higher mortality than males across most study periods. These sex-based differences were statistically significant in Summer 2023 (p = 0.018), Winter 2023 (p = 0.034), and Summer 2024 (p = 0.0004). In contrast, no significant sex difference was observed during Winter 2024 (p = 0.91).

Seasonal variation in mortality was observed exclusively in females when comparing summer periods. Specifically, female mortality in Summer 2024 was significantly higher than in Summer 2023 (p = 0.03), whereas male mortality showed no significant difference between these periods (p = 0.85). Importantly, differences in male and female survival rates are not explained by a positive linear relationship between body size (TL, mm) and survival (Supplementary Materials 3). Across all body sizes, female mortality rates differed clearly from male survival rates (Fig S4).

The model included Site as a random effect to account for potential spatial variation; however, the estimated variance attributed to Site was negligible (variance = 1.7 × 10⁻¹²). Visual inspection of the random effects revealed a normal distribution without notable outliers, indicating that site-specific differences had minimal influence on mortality patterns. This was despite cleaner densities varying by a mean factor of approximately 3.1 between the least and most populated sites. Consequently, we did not pursue alternative models treating site as a fixed effect.

The growth rate of cleaner wrasse varied significantly between seasons (Type II F-Wald Tests: F = 58.59, df = 1, p < 0.001) but also as an interaction between season and year (Type II F-Wald Tests: F = 33.96, df = 1, p < 0.001; Fig 3A). Post hoc analyses revealed that growth rates during the El Niño summer 2024 were significantly lower than in all three other seasons (all p < 0.001). The only other significant difference was that fish grew more slowly in summer 2023 than in winter 2024 (p = 0.005).

**Fig 3.**
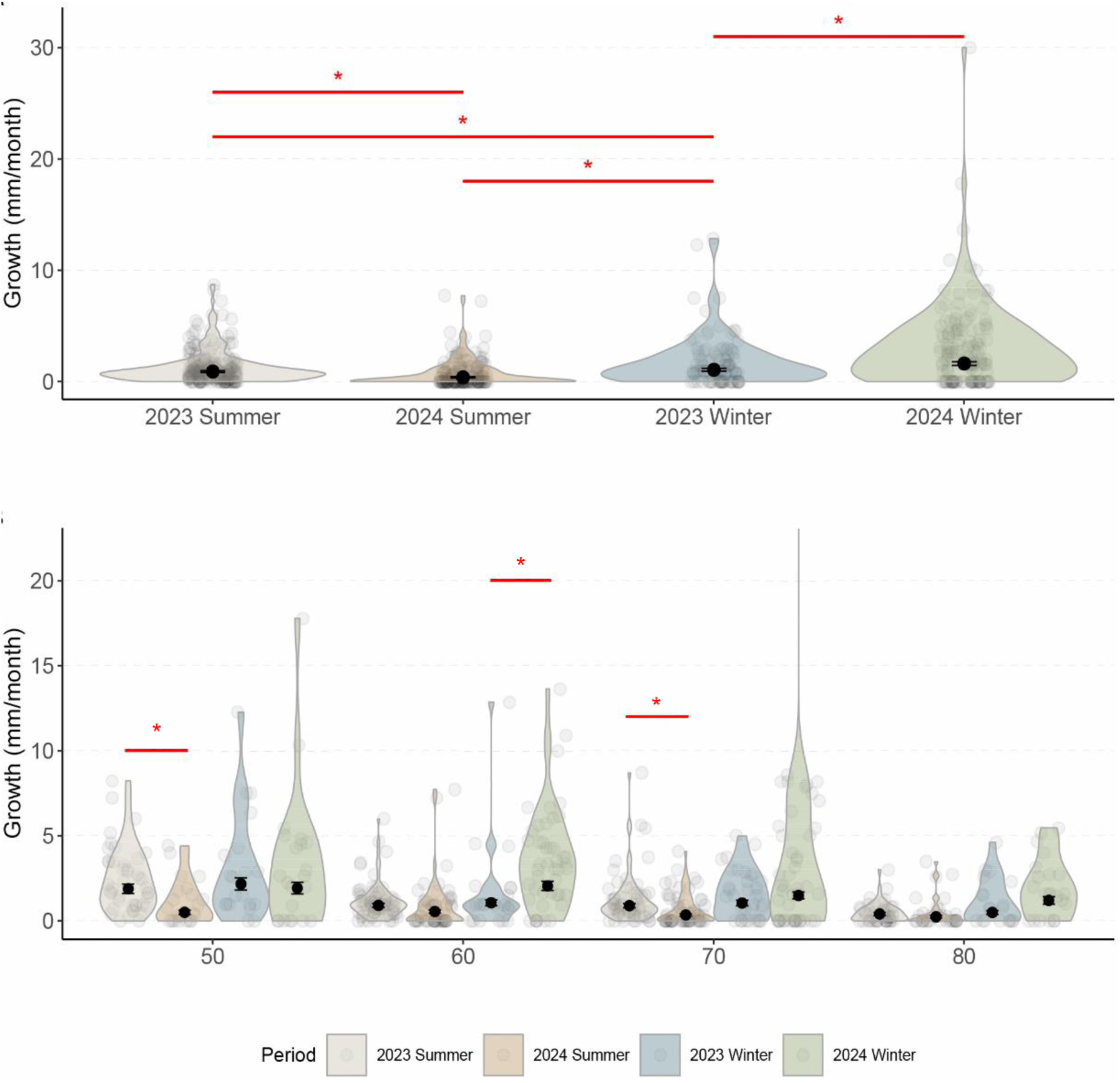
Growth rate variation in cleaner wrasses before, during, and after the 2024 ENSO. Distribution of the cleaner wrasse’s (A), growth rates across time periods, and (B), growth rates across time periods and size classes shown as violins. The jitter represents raw individual-level growth rates, while the overlaid points and error bars represent model-estimated means ± standard errors (SE).

As expected in a species where growth declines with age, growth rates differed significantly across size classes (Type II F-Wald Tests: F = 12.85, df = 3, p < 0.001; Fig 3B). The effect of size class on growth was also year-dependent (Type II F-Wald Tests: F = 4.11, df = 3, p = 0.009). Post hoc analyses indicated that the reduced growth during summer 2024 was most pronounced in individuals from the 50 mm and 70 mm size classes, while the increased growth observed in winter 2024 was primarily driven by individuals in the 60 mm size class.

A second model assessed the role of sex in cleaner wrasse growth rates. This model revealed a significant main effect of sex, with females generally exhibiting faster growth than males (Type II Wald-Chisq Tests: Chisq = 19.8, df = 1, p < 0.001). However, this sex effect varied significantly between years (Type II Wald-Chisq Tests: Chisq = 5.71, df = 1, p = 0.02, Fig 4). Post hoc comparisons revealed a complex pattern. In 2023, a non-bleaching year, females grew significantly more (+45.6 %) during winter compared to summer (p = 0.03), while males did not show a seasonal difference (p = 0.16). When comparing sexes, females grew approximately 44.1% faster than males in summer and 100.2% faster in winter (*p* < 0.0001 for both), indicating a pronounced and seasonally variable sex-based difference in growth patterns under normal environmental conditions. In contrast, in 2024, both males and females exhibited significantly lower growth rates in summer compared to winter (p < 0.0001). Specifically, in summer 2024, the growth rates for females and males were approximately 74% and 61% lower, respectively, compared to winter 2024. Moreover, no significant sex differences were observed in either season (p > 0.1), suggesting that environmental stress during bleaching suppressed growth and erased typical sex-based growth differences, potentially due to shared energetic constraints or altered resource availability. Yearly comparisons revealed opposing trends for males and females. For females, growth in summer 2024 was significantly lower than in summer 2023 (p = 0.0001), while there was no difference between winters (p = 0.07). For males, growth rates were similar across summers (p = 0.997), but growth was significantly higher in winter 2024 compared to winter 2023 (p = 0.01; Fig 4).

**Fig 4.**
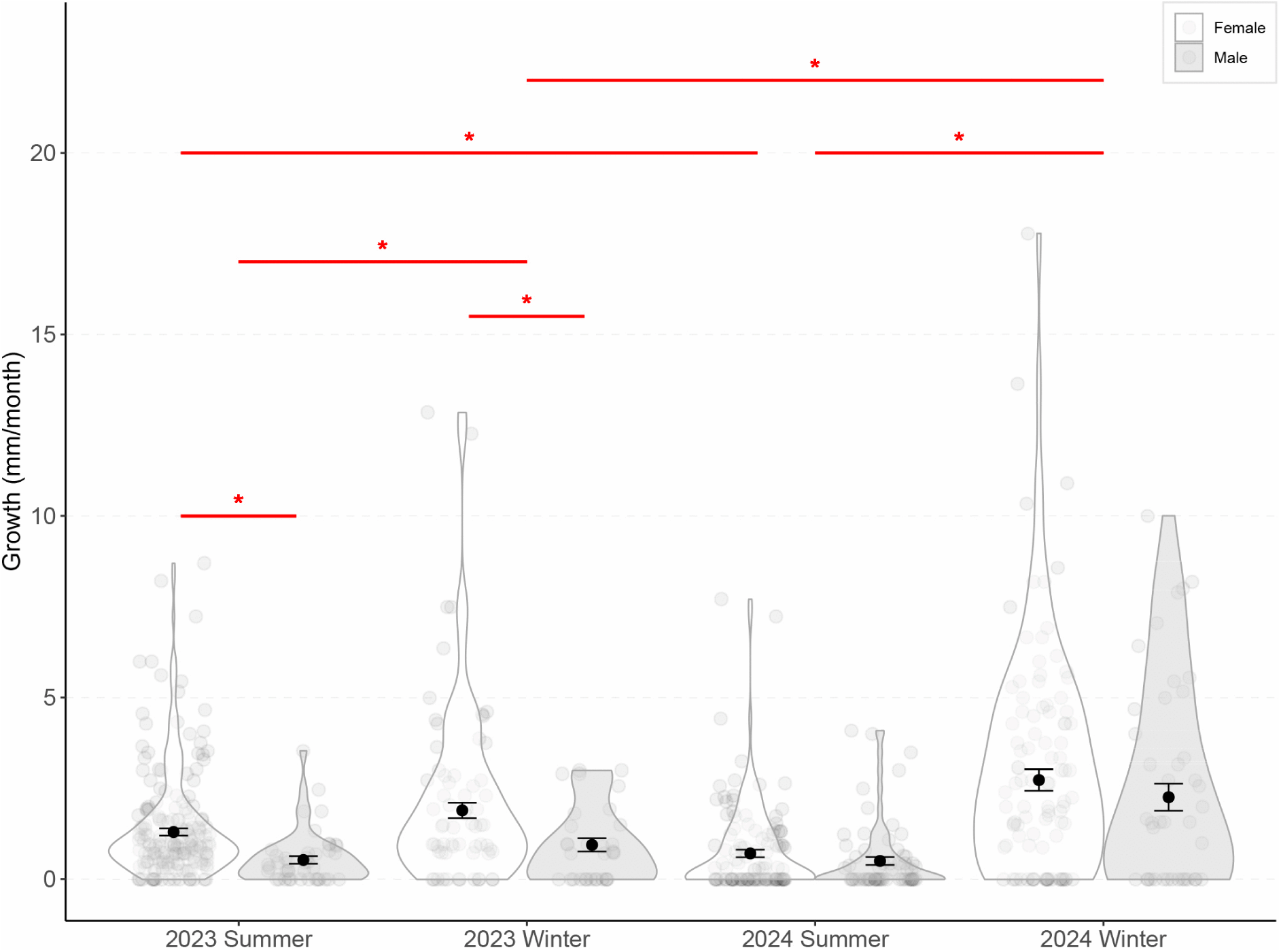
Sex-specific growth rates of cleaner wrasses before, during, and after the 2024 ENSO. Distribution of Male and Female Cleaner wrasse’s growth rates across study periods shown as violins. The jitter represents observed individual-level growth rates, while the overlaid points and error bars represent model-estimated means ± standard errors (SE).

## Discussion

In dioecious polygynous systems, sex-specific differences in life-history traits and associated trade-offs have led to the expectation that males should have higher mortality rates than females do, which becomes even more pronounced during prolonged extreme weather events. Contrary to this assumption, our dataset revealed the opposite pattern in a protogynous coral reef fish, the cleaner wrasse *Labroides dimidiatus*. This is particularly remarkable as sex change theory predicts its evolution when the two sexes differ strongly with respect to the relationship between age and reproductive output, with the terminal sex having higher mean reproductive outputs (Ghiselin 1969; Warner 1988; Munday et al. 2006). Thus, cleaner fish males have not only higher fitness gains but also higher survival rates than females do, both under normal and under extreme weather conditions.

The 2024 ENSO event, marked by intense thermal stress on the Great Barrier Reef (NOAA 2019; Australian Institute of Marine Science 2025), clearly counted as a major disturbance to the cleaner fish population around Lizard Island, as growth was significantly reduced in both sexes compared to other time periods. However, male mortality did not increase, in stark contrast to an about three times increased mortality in females compared to the previous Australian summer. Previous studies on other species suggest that thermal sensitivity can vary across life stages, with younger individuals often showing greater resilience than larger, older ones (Peck et al. 2009; Clark et al. 2013; Peck et al. 2013; Messmer et al. 2017). This has been attributed to a mismatch between oxygen demand and supply at higher temperatures: warming increases metabolic demand while reducing oxygen solubility, which disproportionately affects larger individuals with lower aerobic scope (Peck et al. 2013). By contrast, juveniles tend to have relatively greater aerobic capacity and may better maintain physiological function under thermal stress. Interestingly, the logic does not appear to apply to our study species. Instead, a potential explanation for the increased female mortality during the ENSO event is that they are more vulnerable to direct physiological impacts of elevated temperature (Neuheimer et al. 2011). However, high temperatures not only affect corals and fish populations but also gnathiid isopods, the main ectoparasites of coral reef fishes in the Indo-Pacific (Sikkel et al. 2019). Thus, it is a non-mutually exclusive possibility that cleaner fish faced a reduced amount of food availability, despite the stable populations of other reef fish species (See supplementary analysis 2), and that female cleaner fish suffered more than males from that reduction.

The combination of the two factors is consistent with known effects of warming on metabolic rate: elevated temperatures increase standard metabolic rate (SMR), which, if not offset by higher energy intake (Gillooly et al. 2001), leads to slower growth and higher mortality (Gillooly et al. 2001; Sheridan and Bickford 2011). Nevertheless, the question arises why females are more vulnerable than males in protogynous species. We propose two hypotheses. First, females may consistently have higher energy demands compared to males because they need to invest in egg production for current reproductive success, as well as in growth, to eventually transition into the sex with the higher mean reproductive rate. The inevitable trade-off with survival may have been particularly severe during the ENSO event. As it stands, spawning is rarely observed during focal observations, even in ‘normal’ seasons, preventing us from obtaining quantitative data on its occurrence. Personal observations by the main diver can only confirm that females did not entirely abandon spawning activities during the ENSO. Second, we hypothesize that relatively higher male survival in protogynous species may be due to males being on average of higher quality, as they all survived the challenges associated with the female phase. Future studies should investigate candidate quality components that could differ between the sexes, contrasting physiological features like immune functioning with behavioral and cognitive aspects. Sex differences in cognition have been observed in a variety of species (Geary 1995), including cleaner fish (Triki and Bshary 2021 Jul 14), but are typically linked to niche separation and reproduction (Vinogradov et al. 2025) rather than to survival differences.

Survival during winter 2024 exceeded typical seasonal levels, potentially reflecting delayed benefits of increased per capita access to clients (see supplementary analysis 2) and/or reduced intraspecific aggression following the summer decline in cleaner density. Prior studies on the impacts of multiple cyclones and the 2016 ENSO event reported more severe declines in both cleaner and client densities at Lizard Island compared to 2024 (Triki et al. 2018; Triki and Bshary 2019). Differences in bleaching severity or the absence of cyclone-driven physical reef damage during the 2024 event may help explain this discrepancy. However, our current dataset does not include 2016, limiting the strength of any direct comparison.

Coral reefs are among the most climate-sensitive habitats (Walther et al. 2002; Hughes et al. 2003; Parmesan and Yohe 2003; Hoegh-Guldberg 2009; Hughes et al. 2017; Laufkötter et al. 2020; Malhi et al. 2020; Wernberg et al. 2024), already experiencing marked changes in fish distribution, diversity and community structure due to repeated extreme events (Sylvester 1972; Kim et al. 2001; Jones et al. 2004; Brierley and Kingsford 2009; Eliason et al. 2011; Ferrari et al. 2011; Pearce et al. 2011; Tedesco et al. 2013; Ceccarelli et al. 2024; González-Barrios et al. 2025). Cleaner fish are a keystone species in coral reefs, positively affecting client health (Ros et al. 2011; Triki et al. 2016), growth (Waldie, Blomberg, Cheney, Goldizen, and A. S. Grutter 2011), density and diversity (Bshary 2003; Grutter et al. 2003). But also other protogynous hermaphrodites are of major importance for coral reef functioning (Bonaldo et al. 2014; Kuwamura et al. 2020) or represent commercially important species (Zhou and Gui 2010; Kuwamura et al. 2020). Protogynous species are known to suffer disproportionately from fishing pressure, which targets large individuals (Easter et al. 2020). Thus, pending confirmation from studies on additional species, our results are encouraging in that they suggest males of protogynous species may exhibit high resilience to extreme weather events and climate change in general. Furthermore, sex ratios in protogynous species will very quickly return to equilibrium because all new reproductive individuals will be females and because sex change is socially controlled. More generally, our findings contribute to a growing body of evidence that climate change will not affect all individuals equally (Peck et al. 2009; Clark et al. 2013; Peck et al. 2013; Messmer et al. 2017), emphasizing that species with unusual reproductive strategies may exhibit unexpected demographic responses to environmental stress.

### Ethical note

Our research adheres to the ASAB/ABS Guidelines for the Use of Animals in Research, and the manipulations were approved by the Australian Animal Ethics Committee (AEC; permit numbers CA 2024/06/1865 and CA 2022-04-1601). We acknowledge that catching and marking are stressful events for fish. To minimize stress and handling time, tagging was performed directly underwater at the point of capture, with the fish being kept in a hand net to maintain normal seawater flow. Fish were then released immediately within their home territory, close to the substrate. The entire capture-tag-release process lasted less than 2 minutes per individual, and all fish resumed normal behaviour immediately after release. Additionally, a previous study has shown that cleaner fish can be followed immediately after capture and injection to collect data on intra- and interspecific interactions (Soares et al. 2012). According to this study, there is no evidence that the short manipulation would increase mortality. To minimize the bycatch of non-target species, barrier nets were positioned away from areas with high densities of small reef fishes (e.g., damselfish aggregations near coral heads). Additionally, barrier nets were never left unsupervised. If any non-target species entered the net, they would be immediately released with care.

All subsequent data collection involved only non-invasive methodologies. Growth was measured using an underwater stereo-camera system that required only a brief, close-range pass to film each fish. All other behavioural and life-history data were collected through direct underwater observations, with the diver maintaining a distance of at least 2-3 m, which does not affect their behaviour. Indeed, cleaners often swim closer to the observers, forcing them to retreat and increase distance, so as not to hinder the potential clients from approaching the cleaners.

## Acknowledgements

We thank the staff of the Lizard Island Research Station in Australia, L. Vail, A. Hoggett, R. Car, and A. Davie, for their exceptional support and for fostering an outstanding research environment. We are grateful to Dr. F. Cortesi and the University of Queensland for hosting, and to Dr. A. Green for training in transect data collection. We also thank Amann Marine, Carlos Barbanoj, Diane Berger, Sarah Deventer, Estelle Girord, Corentin Lebet, Anna Viglino, and Arthur Vittet, for their invaluable assistance in the field. We also thank Dr. Radu Slobodeanu for the continuous support and statistical guidance. A special thanks to the inspiring young researchers we met on the island, whose enthusiasm and thoughtful conversations made the work environment both stimulating and welcoming. This work was supported by the Swiss National Science Foundation (grant 310030_192673/1 to RB).

## Author contributions and declaration

All authors contributed to the project design. LP conducted the fieldwork, performed data and statistical analyses, and led manuscript writing. RB supervised the project and contributed to conceptual development and revision of the manuscript.

This manuscript presents original work that has not been published or submitted elsewhere. All authors have approved the final version. We have no conflicts of interest to declare.

