## Supplementary Materials for "Persistent male survival advantage in a protogynous hermaphrodite fish"

#### Content

**Supplementary Materials** ... 1

1. **Supplementary Temperature figure** ... 3

Fig. S1: Lagoon shallow water temperature at Lizard Island for the years 2023 and 2024. ... 3

2. **Supplementary coral and fish analysis** ... 3

**Methods** ... 3

**Results** ... 5

Fig. S2: Proportion of coral bleaching and mortality ... 6

Fig. S3: Fish community response before, during, and after the 2024 ENSO event ... 7

3. **Supplementary survival analysis** ... 8

**Methods** ... 8

**Results** ... 8

Fig. S4: Effect of sex and size on the survival of the cleaner wrasse across study periods. .. 9

4. **Supplementary Island figure** ... 9

Fig. S5: Lizard Island study sites ... 9

5. **Supplementary tagging figure** ... 10

Fig. S6: Illustration of the four locations for VIE ... 10

6. **Supplementary VIE figure** ... 10

Fig. S7: Examples of two individuals that can be recognized without VIE ... 10

7. **Supplementary size error analyses** ... 11

Table S1: Stereo camera system’s error ... 12

Fig. S8: Size error of the software ... 12

8. **Supplementary growth model** ... 13

Fig. S9: Comparison of predicted growth rate between the raw and adjusted dataset ... 14

9. **Supplementary model information** ... 16

Table S2: Glossary for S9.2 Table. ... 16

Table S3: Details about the models. ... 16

**Details for Model 1** ... 17

Table S4: Analysis of Deviance, Type II Chi-Square Tests for Model 1. ... 17

Fig S10: Diagnostic plots for Model 1. ... 17

### 1. Supplementary Temperature figure

**Fig. S1: Lagoon shallow water temperature at Lizard Island for the years 2023 and 2024.** Scatterplot showing temperature in degrees Celsius on the Y axis and year on the X axis

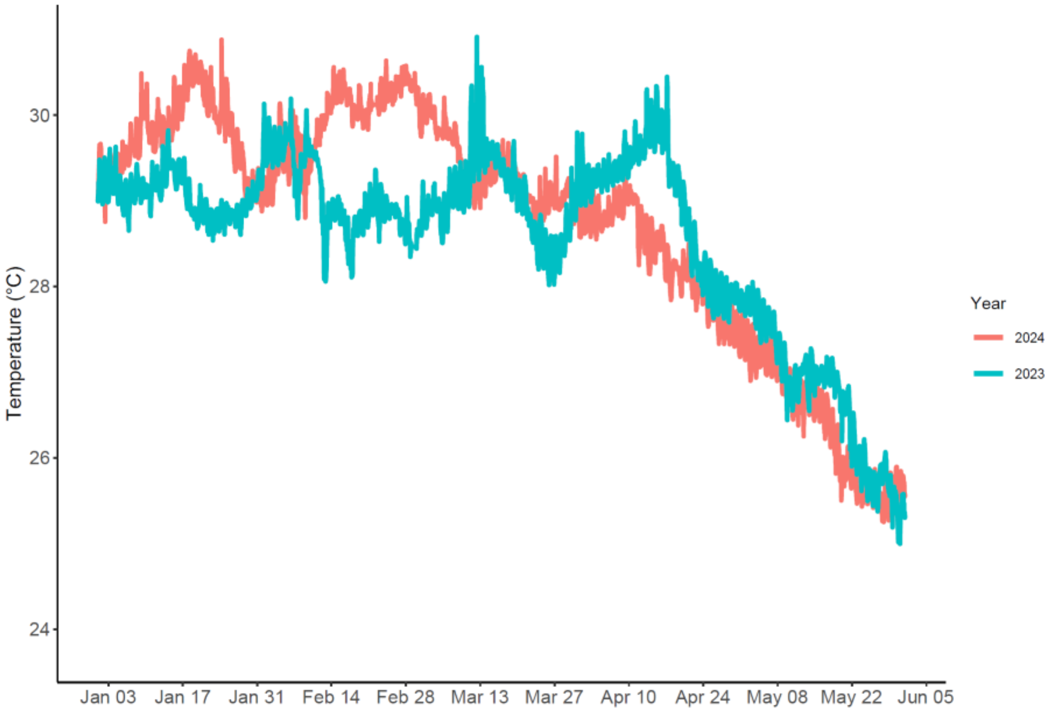

### 2. Supplementary coral and fish analysis

While not central to the core of the manuscript, we regularly collected information on coral bleaching and on fish densities and diversity at our eight study sites. Such information may be of particular interest to colleagues studying climate change, and it provides more context to the conditions that cleaner fish faced during the 2-year study period.

#### Methods

Fish counts included all reef fish, except for nocturnal and cryptic species. Fish censuses were conducted in the Australian summers and winters of 2023 and 2024. Counts were stratified into the two habitats of the reef in which our focal cleaner resided: The reef crest and the reef base.

The reef crest is defined as the seaward edge of the reef flat (Green 1994), and the reef base as the bottom of the reef slope, where it joins the sand flat (Green 1994). Each fish census involved five 30-meter transects, spaced 2 meters apart, running parallel to the reef edge within each habitat for each site. Counts were conducted during three swims: i) the first swim was used to count large fish swimming above the reef within a 5m belt, ii) the second to count medium-sized fish swimming on the reef within a 3m belt, iii) and the last one to count fish species of the family Pomacentridae within a belt of 2m. The 3m belt width for medium-sized fish was selected based on a previous study demonstrating its effectiveness in counting wrasses(Green 1994). Each transect swim was preceded by a 2-minute wait to reduce the impact of diver disturbance from the previous count. Each transect was swum at a constant speed and performed in approximately 10 minutes. Only individuals larger/equal to 4cm were considered in our counts. Fish were identified at the species level, following the WORMS nomenclature, whenever possible. A total of 343 species and 51 families were encountered over the years.

We took advantage of our Fish census methodology to concurrently collect an additional low key dataset on the conditions of corals at our study Sites. A fourth swim was performed along each fish transect to document the reef's condition. Photographs were taken every 2 m using an Olympus TG-6 camera and a 1 $m^2$ quadrat frame, resulting in 15 images per 30 m transect. Photo-quadrats were taken at both the reef crest and reef base habitats at each study site during the austral summers and winters of 2023 and 2024.

To assess bleaching and coral mortality over time, we scored photographic quadrats taken at each site. Each photograph was evaluated independently for the presence of bleaching and coral mortality, resulting in two binary variables indicating whether the respective condition was observed (1) or not (0). These binary observations were aggregated per sampling period and habitat type (reef crest vs. slope) to calculate proportions of photos showing bleaching (Model 5) and coral mortality (Model 6).

Model 4 analyzed the proportion of bleached photos using an LMER with an arcsine-square-root transformation to stabilize variance and approximate normality. Fixed effects included sampling period and habitat, while random intercepts accounted for site-level variation. Variance heterogeneity across sampling years was modelled using a variance-identity structure, allowing residual variance to differ by sampling period and improving model fit.

Model 5 modelled coral mortality as a binomial response using a GLMM. Fixed effects included the sampling period and habitat, while the random effects structure was nested to capture repeated measures across transects, sites, habitats, and Dates.

For client fish densities, we first modelled total fish abundance across years and seasons using an LMER, following a Gaussian distribution (Model 6.1). Fish density was log-transformed to normalize the distribution and stabilize variance, as exploratory data revealed positive skewness. Fixed effects included year, season, and their interaction to capture temporal variability in fish abundance. To account for spatial structures and repeated measures, random intercepts were included for site nested within habitat, reflecting the hierarchical sampling design across the 8 sites.

In Model 6.2, we further examine the differences between small (total length $\leq$ 10 cm) and large clients (total length > 10 cm) using another LMER. Client density was again log-transformed to improve normality. Fixed effects included the sampling period, the client type and their interaction, with the same nested random effects structure as Model 1. Client density was calculated as the number of individuals per $100m^2$.

### Results

From our own data collection, the proportion of pictures of 1 $m^2$ sections containing bleached corals varied significantly across sampling periods (Type II, Wald Chi-square tests: Chisq = 2244.51, df = 2, $p < 0.0001$); S2A Fig). Specifically, bleaching was lower in summer 2023 and significantly increased in all subsequent periods (all $p < 0.0011$), peaking in summer 2024 ($p < 0.0001$). Bleaching was also more pronounced on the reef crest than on the slope (Type II, Wald Chi-square tests: Chisq = 7.6, df = 1, p = 0.006). Coral mortality followed a similar pattern, with the proportion of pictures showing dead coral increasing progressively over time (Type II, Wald Chi-square tests: Chisq = 118.6, df = 2, $p < 0.0001$). However, mortality did not differ significantly between reef crests and slopes (Type II, Chi-square tests: Chisq = 0.29, df = 1, p = 0.59). Post hoc comparisons revealed significant differences between all sampling periods (S2B Fig).

**Fig. S2: Proportion of coral bleaching and mortality.** Proportion of coral bleaching and mortality before, during, and after the 2024 ENSO event. *a*, jittered point data of the proportion of transect photos showing visible coral bleaching. *b*, and hard coral mortality. Overlaid points and error bars represent model-estimated means ± standard error (SE). Individual points represent individual Transects at each study site.

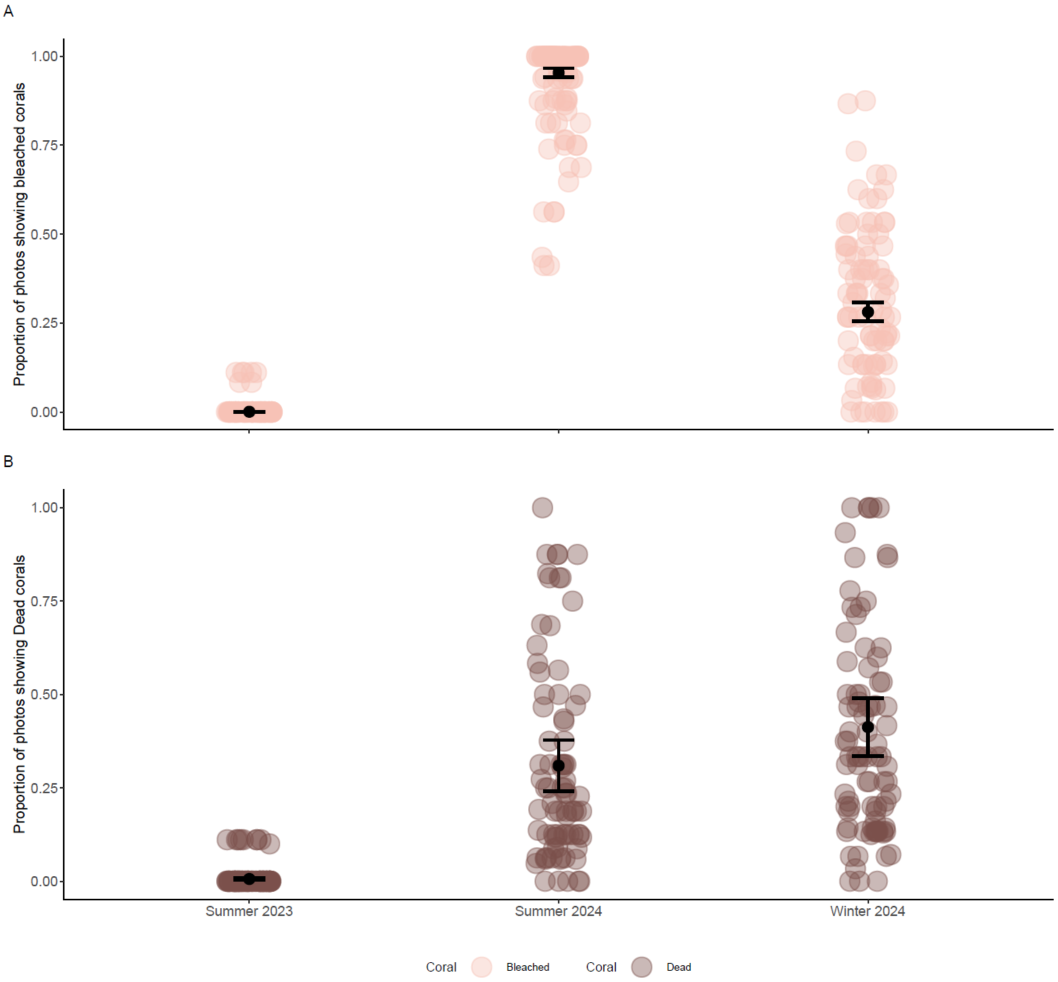

Despite the negative impact on corals, there was no evidence that overall fish populations at our sites experienced increased mortality during the El Niño event compared to the previous year. Instead, total fish density increased markedly from 2023 to 2024 (Type II, F-Wald test: $F = 86.1539$, $df = 1$, $p < 0.0001$), and it was significantly higher in summer than in winter (Type II, F-Wald test: $F = 7.9278$ $df = 1$, $p = 0.005185$), showing no significant interaction between year and season (Analysis of deviance type II F-Wald test: $F = 0.0824$, $df = 1$, $p = 0.774209$; Fig 1A).

Fish density patterns differed between client types with a significant interaction between sampling period and client category (Type II, F-Wald tests: F = 12.719, df = 3, $p < 0.0001$; Fig 1B). This was primarily driven by an increase in the densities of small fish species (<10 cm maximum total length) in both summer and winter 2024 compared to the same seasons in 2023 (all $p < 0.0001$). In contrast, densities of larger fish species (≥ 10 cm max body length) remained stable across years (summer: $p = 0.89$; Winter: $p = 0.99$). Small species were consistently more abundant than large species across all periods (Type II, F-Wald tests, F = 536.92, df = 1, $p < 0.0001$).

**Fig. S3: Fish community response before, during, and after the 2024 ENSO event.** Distribution of (A), general fish density (individuals per 150 m²) and (B), large and small reef fish species density across time periods shown as violins and jitter of the raw transect-level observations at each study site. The overlaid points and error bars represent model-estimated means ± standard errors (SE).

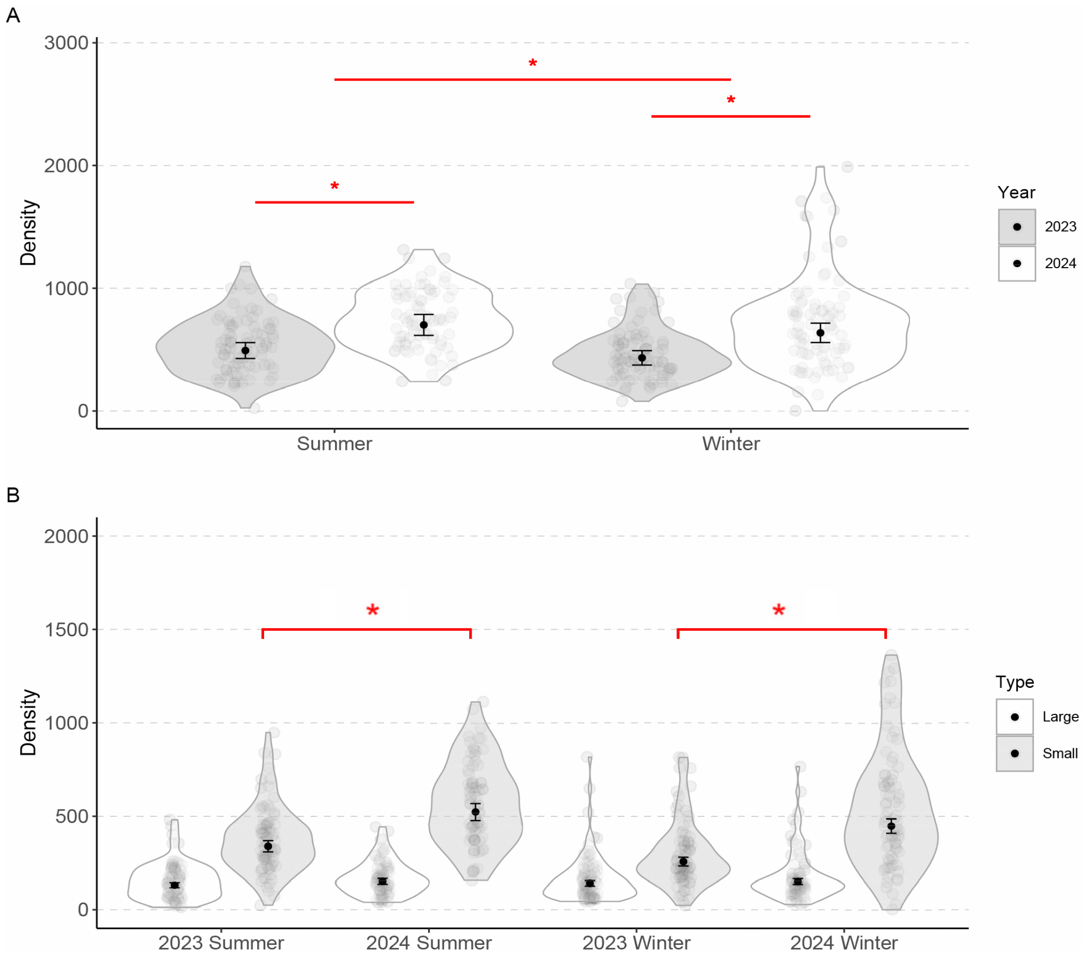

### 3. Supplementary survival analysis

#### Methods

We ran an additional model (Model 7) to test whether the difference in male and female survival could be explained by a simple positive linear relationship between body size (TL, mm) and survival probability. Because males are generally the larger individuals within a group, we aimed to determine whether their higher survival could be attributed to a potential association between larger size and higher survival in cleaner wrasses.

To address this, we combined the presence–absence survival dataset with the size dataset. For each study period, individuals were assigned a survival score of 1 (survived) or 0 (did not survive), and their initial body size was included. Only individuals with recorded size measurements were used in this analysis.

We fitted a generalized linear model (GLM) with a binomial error distribution, using survival (0/1) as the response variable and size, sex, and their interaction as explanatory variables.

#### Results

The only significant predictor was sex (Analysis of Deviance, Type II test: Chisq = 26.79, df = 1, $p < 0.0001$). Body size was not significant (Chisq = 0.0002, df = 1, $p = 0.99$) and did not influence the effect of sex (Chisq = 0.07, df = 1, $p = 0.786$; S3 Fig).

**Fig. S4: Effect of sex and size on the survival of the cleaner wrasse across study periods.** Model predictions are shown for both males (blue) and females (red) across study periods.

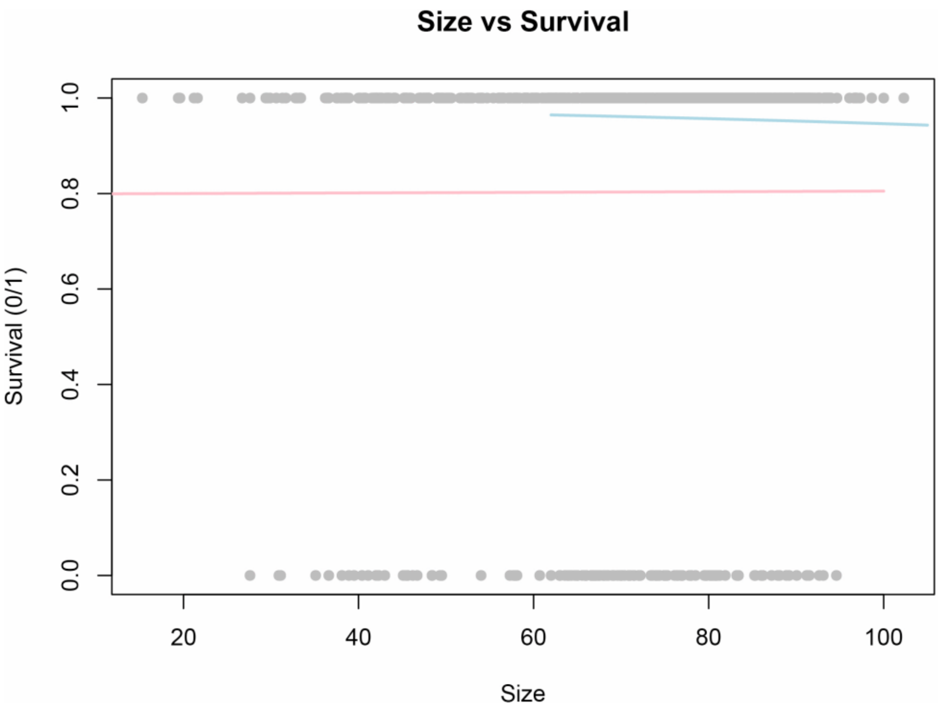

### 4. Supplementary Island figure

**Fig. S5: Lizard Island study sites.** Grey represents the land, and blue dots represent the reef sites studied.

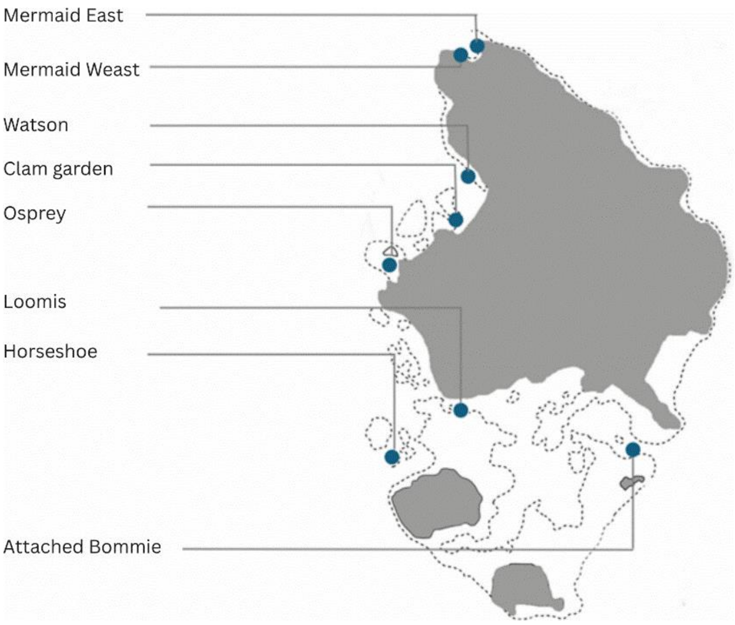

### 5. Supplementary tagging figure

**Fig. S6: Illustration of the four locations for VIE.** Two dots are shown at each location, since colors can be injected in sequence in the same area.

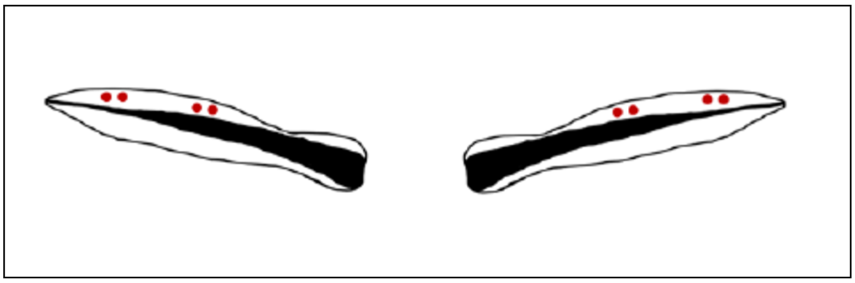

### 6. Supplementary VIE figure

**Fig. S7: Examples of two individuals that can be recognized without VIE.** On the left, the fish has a natural black marking on its beige body band. On the right, the fish shows a natural white interruption of the black band.

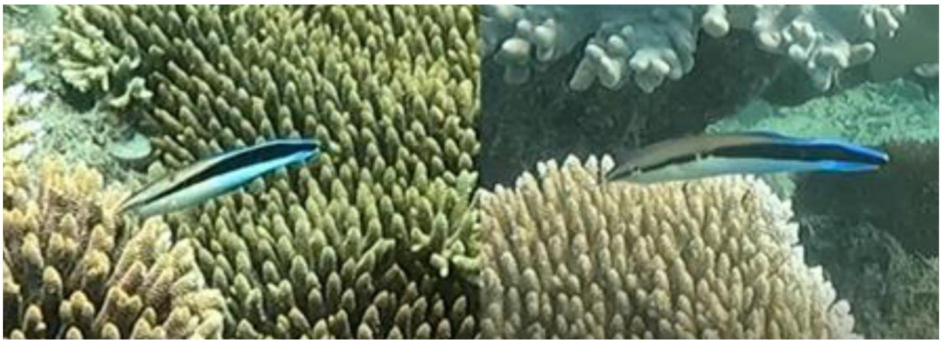

### 7. Supplementary size error analyses

We assessed the growth of each focal fish using a calibrated underwater stereo-photogrammetric system, consisting of two GoPro Hero 8 cameras mounted on a fixed rig (Seager 2006). Footage was processed with EventMeasure (Seager 2006) which enables precise three-dimensional measurements based on synchronized stereo images. Before data collection, the system was calibrated using CAL, a dedicated software that ensures accurate geometric alignment between the camera pair (Seager 2006). This stereo setup significantly improves measurement precision over traditional visual estimations (Michael et al. 2011). In the first three seasons, size measurements were taken every two weeks, whereas in winter 2024, they were collected at a reduced frequency of once a month.

The error of this software is known to be around 1-2mm (Euan et al. 2010). To further assess the accuracy of the stereo camera system for our study, a calibration bar with three known distances was used to measure the system's error. Measurement error was ± 1.13mm using the calibration bar and ± 1.81mm of wild cleaner fish due to movement and growth between measurements (S7 Fig, S7 Table).

Because measuring a moving object is more challenging, the software's accuracy was then investigated by comparing the stereo camera measurements of fish longer than 70cm with manual size measurements taken less than 30 days prior. This method provided an average error of ± 1.81mm. Attempts were made to obtain size measurements using the stereo system less than one week after the fish were manually measured post-capture. However, this proved challenging as the fish required more time to re-acclimate to the presence of humans and often swam too quickly or attempted to escape, making accurate measurements difficult within this short time frame. Nevertheless, errors associated with our fish measurements did not show a significant increase in variance (variance = 2.324) compared to the error observed with the calibration toolbar (variance = 2.264). This suggests that the software's measurement error remains consistent when applied to real fish. The positive shift in the median error for the fish measurements (median = 1.48 mm) reflects the natural growth of the fish over the 30-day period between the two measurements. This consistency in error variance across both methods indicates that the software performs reliably for measuring wild fish.

**Table S1: Stereo camera system's error.** Error in (mm) of the stereo camera system obtained using tool bars of different sizes (small, medium, large), with measurements of fish in the water (Fish), and as an overall mean from the various tool bars.

| Method | Mean Error (mm) |
|---|---|
| Small | ± 0.981 |
| Medium | ± 1.08 |
| Large | ± 1.37 |
| General Tool | ± 1.13 |
| Fish | ± 1.81 |

**Fig. S8: Size error of the software.** The size error of the software obtained using size measurements of focal cleaners ("Cleanerfish") obtained 30 days from their manual sizing, and using the objects of known size ("Calibration"). Boxplots show medians (center line), interquartile ranges (boxes), and 95% data range.

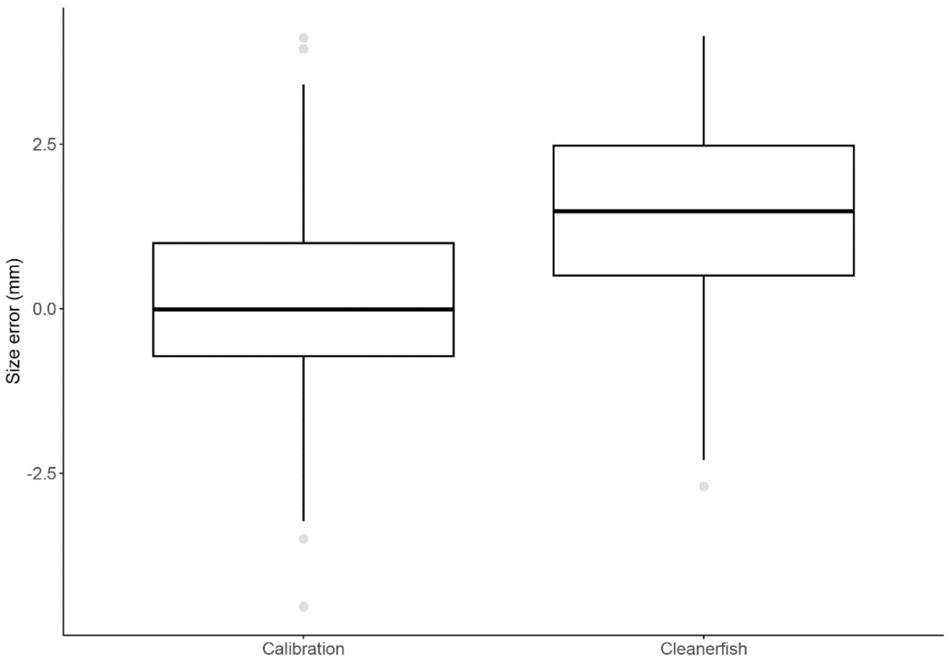

### 8. Supplementary growth model

To evaluate if this correction affected our interpretation of growth dynamics, we ran an additional model (Model 3.3) using the unadjusted, raw growth data. This model was not used for hypothesis testing, but rather to extract estimates and prediction distributions, verifying that adjusting negative values in the main models did not introduce substantial bias. Because this exploratory model was intended for comparisons rather than inference, strict adherence to model assumptions was not required and was not entirely achievable given the structure of the raw data. This supplementary model was an LMER using the logarithm of growth as the response and year, sex, and season as fixed effects. Random intercepts were included for individual ID nested within site to account for repeated measures and site-level variation. Each observation was weighted by the inverse of the squared daily measurement error ($1/DE^2$), implemented through a fixed variance structure (varFixed), to give greater weight to more precise measurements. Additionally, we included a variance identity function (varIdent) to allow residual variances to differ across levels of the grouping variable (including year, sex and season). These two variance structures were combined using varComb() to simultaneously address both observation-level and group-level heteroscedasticity.

The error associated with growth measurements obtained from the sizing software was calculated by combining the errors of the initial and final size measurements (attributed to be ± 0.981, the error calculated for small lengths). Specifically, growth error was computed as the square root of the sum of the squared size errors at both time points using equation (2).

$$\text{growth error} = \sqrt{(\text{size error}_{\text{initial}})^2 + (\text{size error}_{\text{final}})^2} \qquad (2)$$

This growth error was then standardized dividing it by the number of days between measurements to account for varying measurement intervals, resulting in the daily error (DE).

**Fig. S9: Comparison of predicted growth rate between the raw and adjusted dataset.** The (A), Predicted growth rates for female and (B), male cleaner fish across years and seasons, based on models using raw (with negative values) vs adjusted (negative values were adjusted) growth data. Points show median predictions, and error bars show 95% confidence intervals.

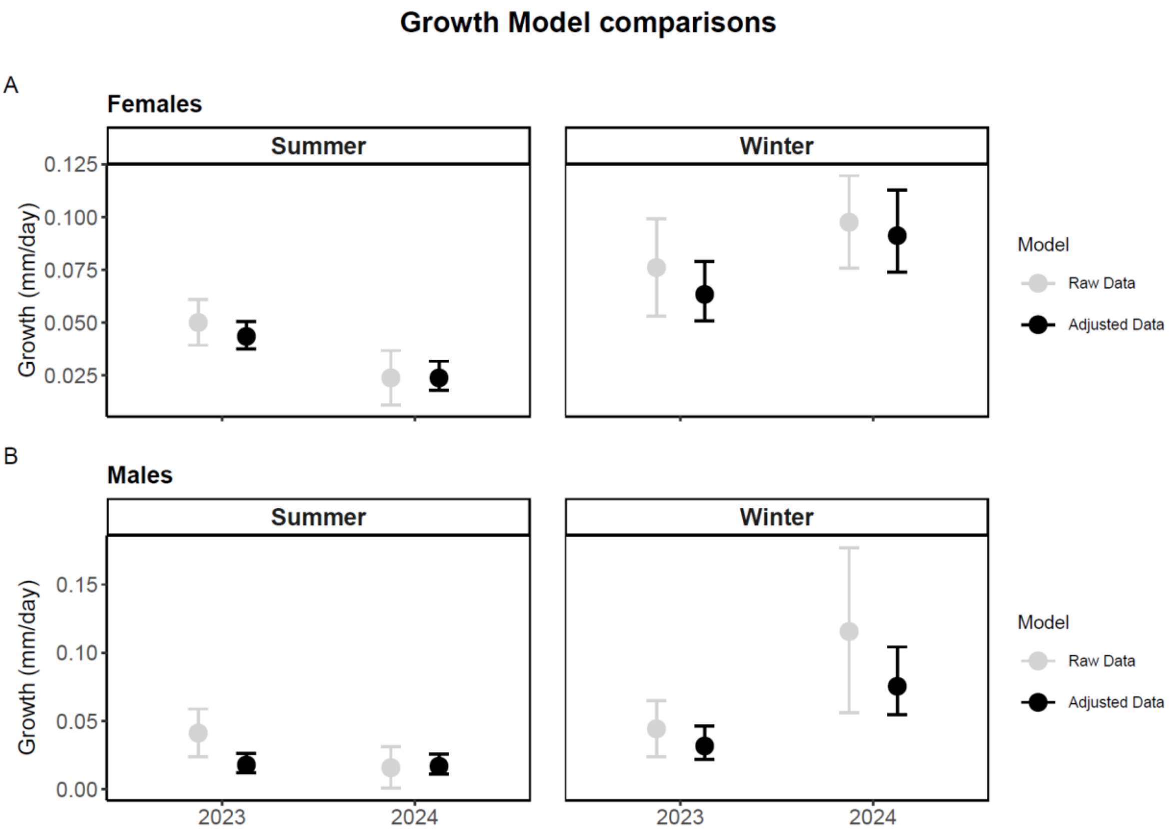
Growth Model comparisons
A
Females
Summer
Winter
Growth (mm/day)
0.125
0.100
0.075
0.050
0.025
Model
Raw Data
Adjusted Data
B
Males
Summer
Winter
Growth (mm/day)
0.15
0.10
0.05
0.00
2023
2024
Model
Raw Data
Adjusted Data

### 9. Supplementary model information

**Table S2: Glossary for Table S3.**

| Term | Definition |
|---|---|
| Class | Size class (TL: 60-69mm, 70-79mm, 80-90mm) |
| DE | Error of the Sizing Software (see Daily error in the method section) |
| Density | Adult cleaner fish reef density within a deme, standardized by $100m^2$ |
| Fish-Density | Client densities counted during transects standardized per $100m^2$ |
| Growth | Growth rate (mm/day) |
| ID | Cleaner ID used as a random factor to control for repeated measures |
| | Summer 2023, summer 2024, winter 2023, and winter 2024. |
| Type | Large or small clients |
| Season | Summer or winter |
| Sex | Male or Female |
| Habitat | Reef Crest or Slope. |

**Table S3: Details about the models.**

| Model | Type | Family(link) | Sample Size | Formula |
|---|---|---|---|---|
| **1** | LMER | Gaussian | 32 obs | log(Density+2) ~ Season * Year +(1\|Site) |
| **2.1** | LMER | Gaussian | 617 obs<br>402 ID | log(growth+0.01)~(Class+Season)*Year+(1\|Site)+(1\|ID) |
| **2.2** | GLMER | Tweedie(log) | 621 obs<br>406 ID<br>447 Females<br>174 Males | growth~ Sex * Year *Season +(1\|ID) + (1\|Site), dispformula =~0+Year |
| **2.3** | LME | Gausian | 621 obs<br>406 ID<br>447 Females<br>174 Males | Log(growth+1.61) ~ Sex * Year * Season, random = ~1\|Cleaner_ID/Site, weights = varComb( varFixed(~I\|DE^2)), WarTdent(form =~1\|mix) |
| **3** | LME | Gaussian | 65 obs<br>33 Females<br>32 Males | asin(sqrt(Mortality)) ~ Period * Sex, random = ~1\|Site, weight=varIdent(form=~1\|Period) |
| **4** | LMER | Gaussian | 260 obs | asin(sqrt(bleached/total)) ~ MY + Habitat, random = ~1\|Site, weights = varIdent(form=~1\|MY) |
| **5** | GLMM | binomial | 260 obs | cbind(dead, tot) ~ MY + Habitat + (Transect/Site/Habitat/Date) |
| **6.1** | LMER | Gaussian | 324 obs | Log(Fish Density+160) ~ Year * Season +(1\| Habitat/Site) |
| **6.2** | LMER | Gaussian | 646 obs | Log(Fish Density+40) ~ Period * Type +(1\|Habitat/Site) |
| **7** | GLM | binomial | 838 obs<br>219 Males<br>619 Females | Survival~Size*Sex |

### Details for Model 1

**Table S4: Analysis of Deviance, Type II Chi-Square Tests for Model 1.**

| Term | Chisq | Df | Pr(>F) |
|---|---|---|---|
| Season | 0.3428 | 1 | 0.558205 |
| Year | 8.3222 | 1 | 0.003916 ** |
| Season:Year | 0.2489 | 1 | 0.617821 |

**Fig S10: Diagnostic plots for Model 1.**

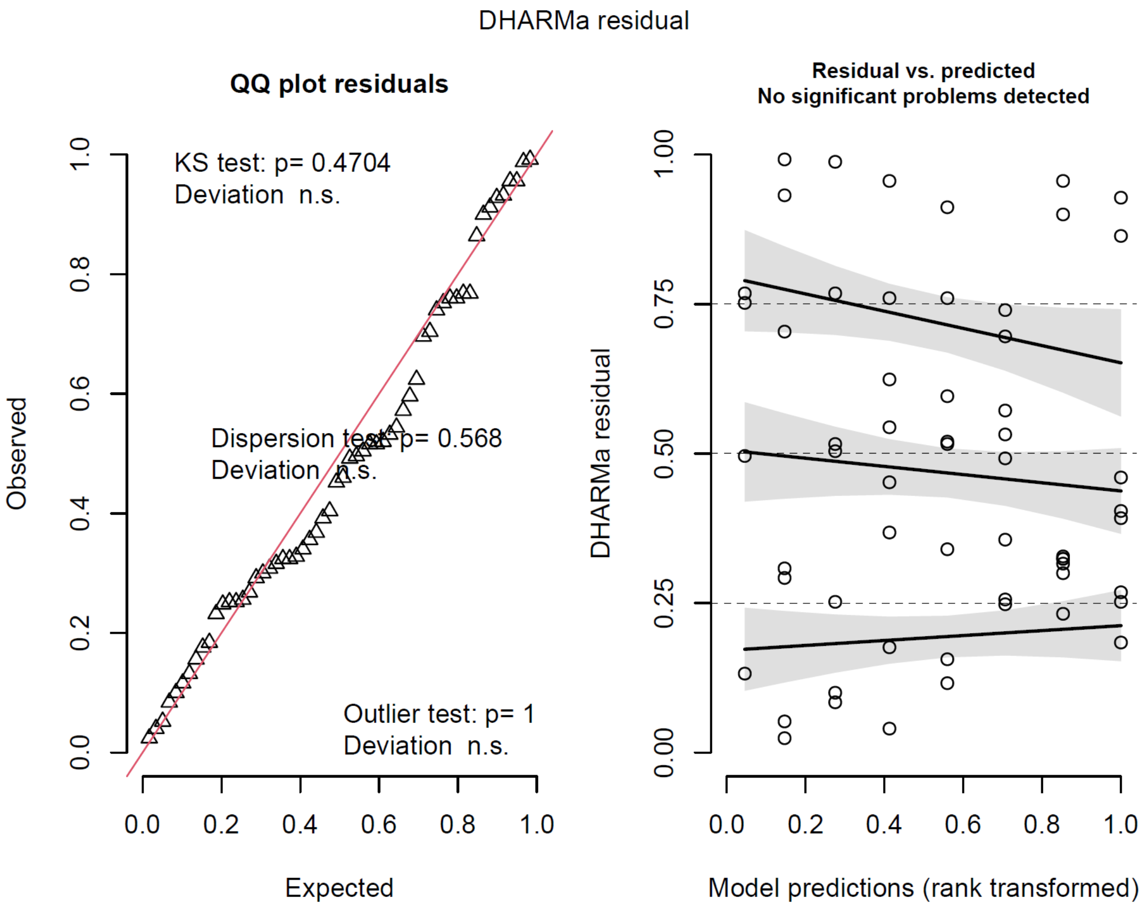

### Details for Model 2.1

**Table S5: Analysis of Deviance, Type II Wald F Tests for Model 2.1.**

| Term | F | Df | Pr(>F) |
|---|---|---|---|
| Class | 12.4499 | 3 | 1.018-07 *** |
| Season | 60.0548 | 1 | 5.78e-14 *** |
| Year | 7.7467 | 1 | 0.005570 ** |
| Class:Year | 3.9789 | 3 | 0.008031 ** |
| Season:Year | 34.9361 | 1 | 6.450e-09 *** |

**Table S6: Emmeans Contrasts by Class for Model 2.1**

| Contrast | Estimate | SE | Df | t.ratio | p-value |
|---|---|---|---|---|---|
| **Class = 50** | | | | | |
| Summer 2023 – Winter 2023 | -0.009370 | 0.01950 | 368 | -0.482 | 0.9631 |
| Summer 2023 – Summer 2024 | 0.040703 | 0.01190 | 368 | 3.419 | 0.0039 |
| Summer 2023 – Winter 2024 | 0.015277 | 0.01540 | 368 | 0.993 | 0.7534 |
| Winter 2023 – Summer 2024 | 0.050073 | 0.01780 | 469 | 2.811 | 0.0263 |
| Winter 2023 – Winter 2024 | 0.024646 | 0.02030 | 446 | 1.214 | 0.6186 |
| Summer 2024 – Winter 2024 | -0.025426 | 0.01300 | 446 | -1.954 | 0.2073 |
| **Class = 60** | | | | | |
| Summer 2023 – Winter 2023 | 0.000274 | 0.00867 | 196 | 0.032 | 1.0000 |
| Summer 2023 – Summer 2024 | 0.016997 | 0.00534 | 196 | 3.183 | 0.0091 |
| Summer 2023 – Winter 2024 | -0.050370 | 0.01350 | 196 | -3.729 | 0.0014 |
| Winter 2023 – Summer 2024 | 0.016724 | 0.00822 | 285 | 2.036 | 0.1774 |
| Winter 2023 – Winter 2024 | -0.050644 | 0.01480 | 271 | -3.421 | 0.0040 |
| Summer 2024 – Winter 2024 | -0.067368 | 0.01310 | 271 | -5.156 | <.0001 |
| **Class = 70** | | | | | |
| Summer 2023 – Winter 2023 | -0.004397 | 0.00813 | 244 | -0.541 | 0.9490 |
| Summer 2023 – Summer 2024 | 0.017896 | 0.00522 | 244 | 3.425 | 0.0040 |
| Summer 2023 – Winter 2024 | -0.016719 | 0.00896 | 244 | -1.865 | 0.2458 |
| Winter 2023 – Summer 2024 | 0.022294 | 0.00740 | 262 | 3.011 | 0.0151 |
| Winter 2023 – Winter 2024 | -0.012321 | 0.01040 | 321 | -1.188 | 0.6347 |

| | | | | | |
|---|---|---|---|---|---|
| Summer 2024 – Winter 2024 | -0.034615 | 0.00812 | 262 | -4.265 | 0.0002 |
| **Class = 80** | | | | | |
| Summer 2023 – Winter 2023 | -0.008854 | 0.00693 | 336 | -1.277 | 0.5783 |
| Summer 2023 – Summer 2024 | 0.003945 | 0.00418 | 336 | 0.944 | 0.7809 |
| Summer 2023 – Winter 2024 | -0.027795 | 0.01020 | 336 | -2.726 | 0.0339 |
| Winter 2023 – Summer 2024 | 0.012799 | 0.00699 | 423 | 1.831 | 0.2602 |
| Winter 2023 – Winter 2024 | -0.018942 | 0.01160 | 490 | -1.636 | 0.3593 |
| Summer 2024 – Winter 2024 | -0.031741 | 0.01010 | 423 | -3.149 | 0.0095 |

**Fig S11: Diagnostic plots for Model 2.1.**

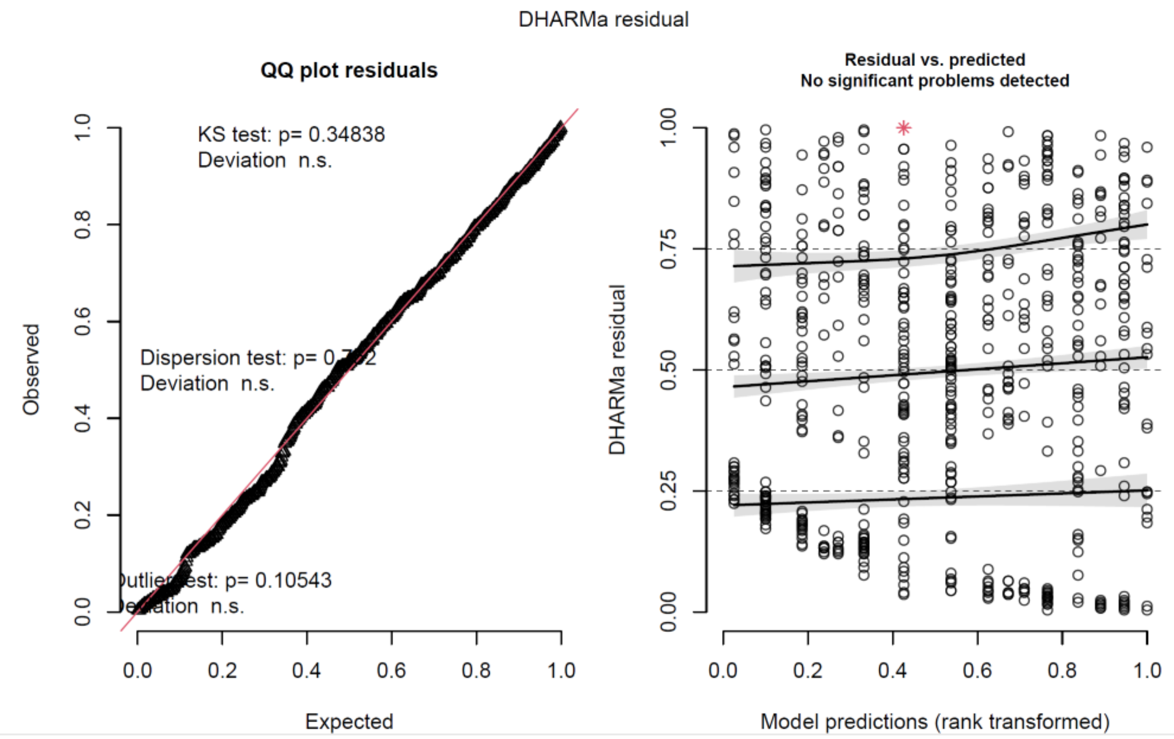

### Details for Model 2.2

**Table S7: Analysis of Deviance, Chi-square Tests for Model 2.2.**

| Term | Chisq | Df | Pr(>F) |
|---|---|---|---|
| Sex | 19.7634 | 1 | 8.764e-06 *** |
| Year | 0.3071 | 1 | 0.57948 |
| Season | 79.3319 | 1 | < 2.2e-16 *** |
| Sex:Year | 5.7258 | 1 | 0.01672 * |
| Sex:Season | 0.7057 | 1 | 0.40088 |
| Year:Season | 27.2558 | 1 | 1.1782e-07 *** |
| Sex:Year:Season | 0.0113 | 1 | 0.91549 |

**Table S8: Emmeans Contrasts by Year for Model 2.2.**

| Contrast | estimate | SE | z.ratio | p-value |
|---|---|---|---|---|
| **Year = 2023** | | | | |
| Female Summer - Male Summer | 0.02568 | 0.00477 | 5.380 | <.0001 |
| Female Summer - Female Winter | -0.01986 | 0.00728 | -2.729 | 0.0322 |
| Female Summer - Male Winter | 0.01186 | 0.00678 | 1.748 | 0.2990 |
| Male Summer - Female Winter | -0.04553 | 0.00788 | 5.777 | <.0001 |
| Male Summer - Male Winter | -0.01382 | 0.00669 | 2.066 | 0.1642 |
| Female Winter - Male Winter | 0.03171 | 0.00934 | 3.396 | 0.0038 |
| **Year=2024** | | | | |
| Female Summer - Male Summer | 0.00698 | 0.00493 | 1.416 | 0.4892 |
| Female Summer - Female Winter | -0.06747 | 0.01010 | -6.688 | <.0001 |
| Female Summer - Male Winter | -0.04945 | 0.01270 | -3.895 | 0.0006 |
| Male Summer - Female Winter | -0.07445 | 0.01040 | -7.180 | <.0001 |
| Male Summer - Male Winter | -0.05643 | 0.01250 | -4.508 | <.0001 |
| Female Winter - Male Winter | 0.01801 | 0.01540 | 1.167 | 0.6479 |

**Table S9: Emmeans Contrasts by sex for Model 2.2.**

| Contrast | Estimate | SE | z.ratio | p-value |
|---|---|---|---|---|
| **Sex = Female** | | | | |
| 2023 Summer - 2024 Summer | 0.019637 | 0.00454 | 4.330 | 0.0001 |
| 2023 Summer - 2023 Winter | -0.019855 | 0.00728 | -2.729 | 0.0322 |
| 2023 Summer - 2024 Winter | -0.047829 | 0.01010 | -4.747 | <.0001 |
| 2024 Summer - 2023 Winter | -0.039492 | 0.00759 | -5.206 | <.0001 |
| 2024 Summer - 2024 Winter | -0.067466 | 0.01010 | -0.04155 | <.0001 |
| 2023 Winter - 2024 Winter | -0.027974 | 0.01150 | -2.424 | 0.0725 |
| **Sex = Male** | | | | |
| 2023 Summer - 2024 Summer | 0.000938 | 0.00489 | 0.192 | 0.9975 |
| 2023 Summer - 2023 Winter | -0.013822 | 0.00669 | 0.00336 | 0.1642 |
| 2023 Summer - 2024 Winter | -0.055496 | 0.01260 | -0.02306 | 0.0001 |
| 2024 Summer - 2023 Winter | -0.014760 | 0.00687 | 0.00290 | 0.0001 |
| 2024 Summer - 2024 Winter | -0.056434 | 0.01250 | -0.02427 | <.0001 |

**Fig S12: Diagnostic plots for Model 2.2.**

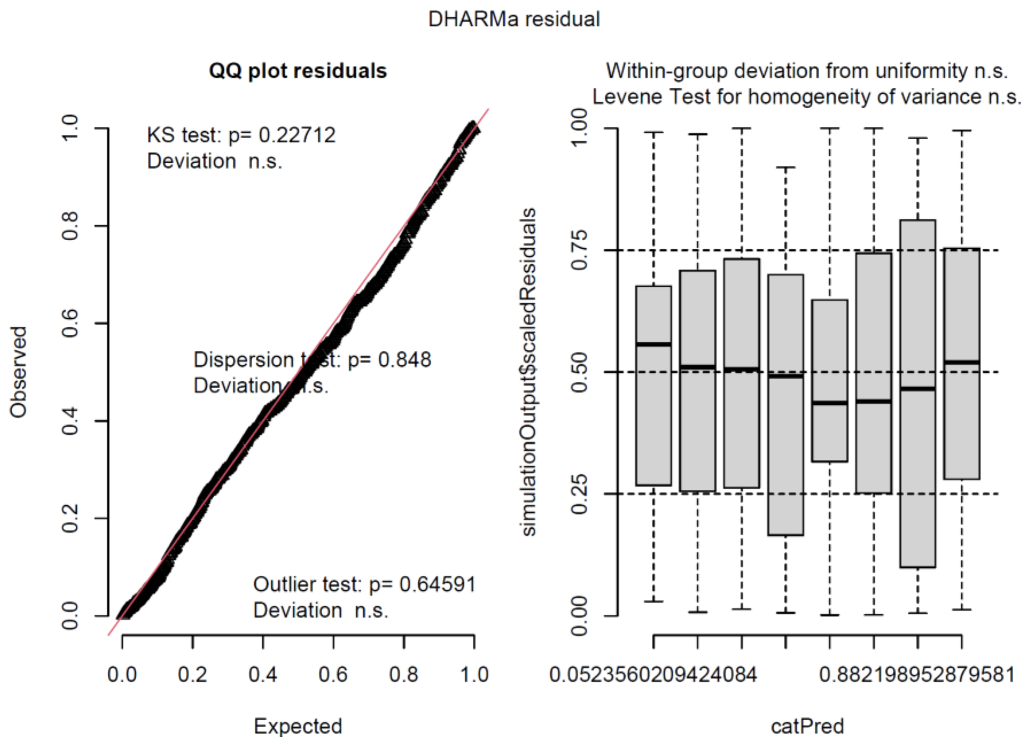

### Details for Model 3

**Table S10: Analysis of Deviance, Chi-square Tests for Model 3.**

| Term | Chisq | Df | Pr(>F) |
|---|---|---|---|
| MY | 8.0835 | 3 | 0.04432* |
| Sex | 17.3317 | 1 | 3.139-e05*** |
| MY:Sex | 7.5822 | 3 | 0.05548 . |

**Table S11: Emmeans Contrasts by Period for Model 3.**

| Contrast | | Estimate | SE | df | t-ratio | p-value |
|---|---|---|---|---|---|---|
| 2023 Summer | Female - Male | 0.1402 | 0.0574 | 50 | 2.443 | 0.0181 |
| 2023 Winter | Female - Male | 0.2188 | 0.1020 | 50 | 2.138 | 0.0374 |
| 2024 Summer | Female - Male | 0.2676 | 0.0706 | 50 | 3.789 | 0.0004 |
| 2024 Winter | Female - Male | 0.0079 | 0.0682 | 50 | 0.116 | 0.9083 |

**Table S12: Emmeans Contrasts by Sex for Model 3.**

| Contrast | estimate | SE | df | t.ratio | p-value |
|---|---|---|---|---|---|
| **Sex = Female** | | | | | |
| Summer 2023 – Winter 2023 | -0.1117 | 0.0824 | 50 | -1.1356 | 0.5325 |
| Summer 2023 - Summer 2024 | -0.1795 | 0.0636 | 50 | -2.823 | 0.033 |
| Summer 2023 – Winter 2024 | 0.0623 | 0.0622 | 50 | 1.001 | 0.7492 |
| Winter 2023 – Summer 2024 | -0.0678 | 0.0879 | 50 | -0.772 | 0.8668 |
| Winter 2023 – Winter 2024 | 0.1740 | 0.0869 | 50 | 2.001 | 0.2012 |
| Summer 2024 – Winter 2024 | 0.2418 | 0.0694 | 50 | 3.484 | 0.0056 |
| **Sex = Male** | | | | | |
| Summer 2023 – Winter 2023 | -0.0330 | 0.0835 | 50 | -0.396 | 0.9788 |
| Summer 2023 – Summer 2024 | -0.0521 | 0.0651 | 50 | -0.800 | 0.8542 |
| Summer 2023 – Winter 2024 | -0.0699 | 0.0638 | 50 | -1.096 | 0.6934 |
| Winter 2023 – Summer 2024 | -0.0190 | 0.0879 | 50 | -0.216 | 0.9964 |
| Winter 2023 – Winter 2024 | -0.0369 | 0.0869 | 50 | -0.424 | 0.9741 |
| Summer 2024 – Winter 2024 | -0.0179 | 0.0694 | 50 | -0.257 | 0.9939 |

**Fig S13: Diagnostic plots for Model 3.**

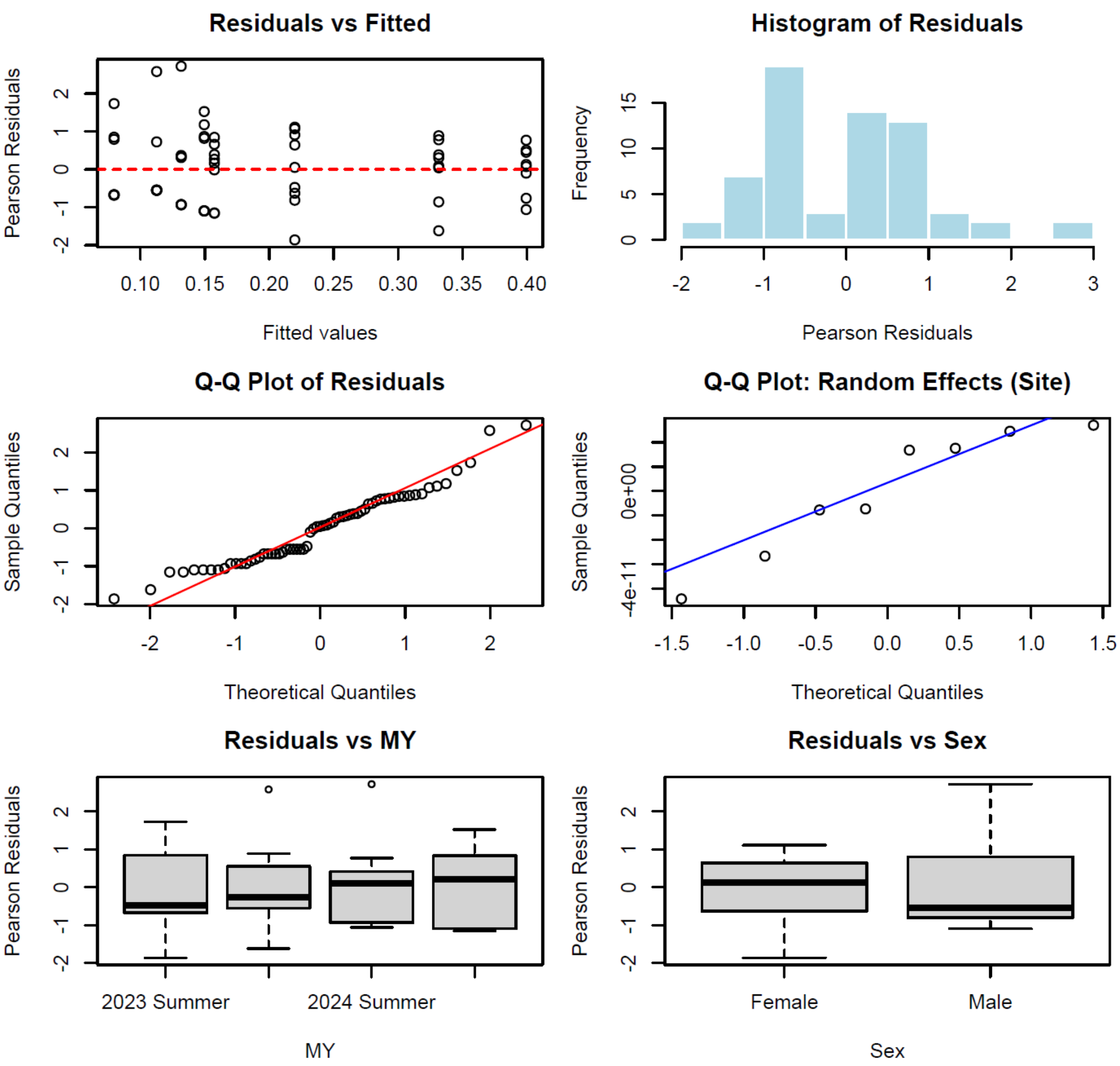

### Details for Model 4

**Table S13: Analysis of Deviance, Type II Chi-Square Tests for Model 4.**

| Term | Chisq | Df | Pr(>F) |
|---|---|---|---|
| MY | 2244.5070 | 2 | < 2.2e-16 *** |
| Habitat | 7.5992 | 1 | 0.005839 ** |

**Table S14: Emmeans Contrasts for Model 4.**

| | Estimate | SE | Df | T ratio | P value |
|---|---|---|---|---|---|
| Summer-2023-Summer 2024 | -1.327 | 0.0294 | 249 | -45.096 | <.0001 |
| Summer 2023-Winter 2024 | -0.5329 | 0.0285 | 249 | -18.756 | <.0001 |
| Summer 2024-Winter 2024 | 0.7945 | 0.0386 | 249 | 20.430 | <.0001 |

Results averaged over the levels of Scenario, Degrees of freedom method : Kenward-roger. Results are given on the log scale (not the response one).

**Fig S14: Diagnostic plots for Model 4.**

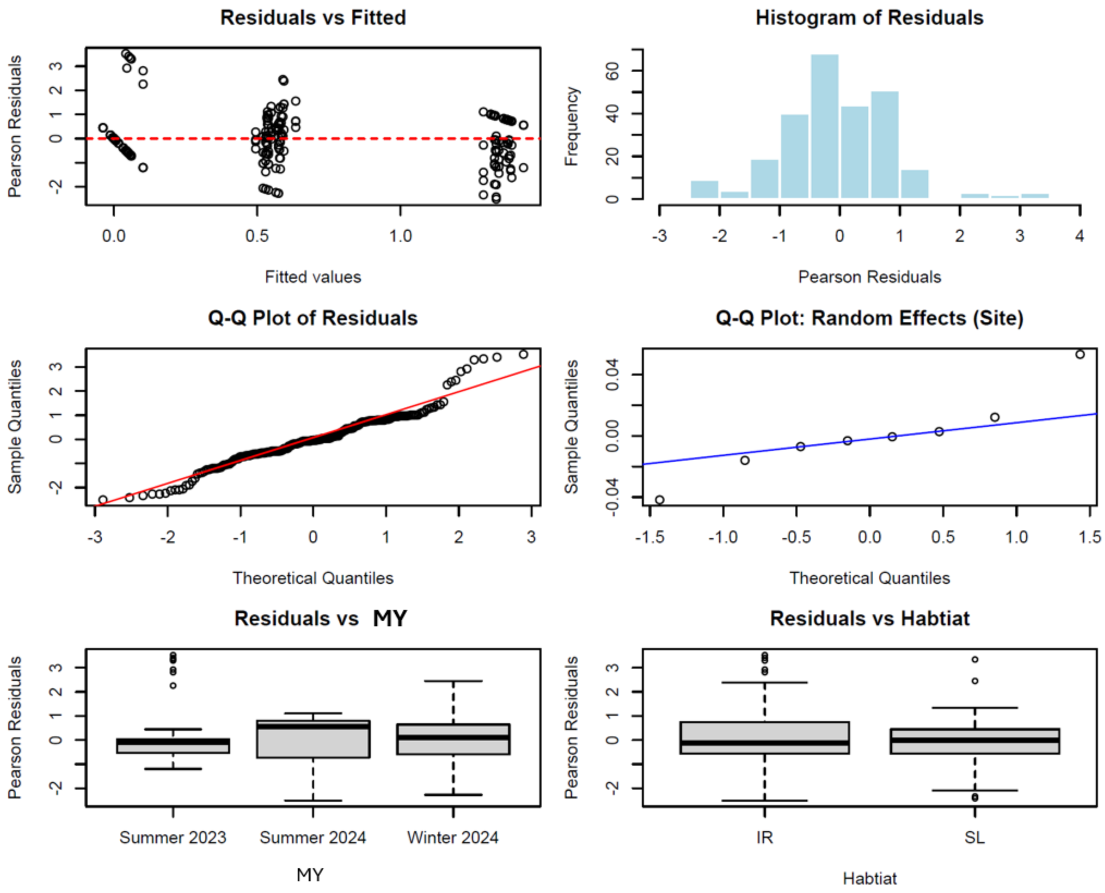

### Details for Model 5

**Table S15: Analysis of Deviance, Type II Chi-Square Tests for Model 5.**

| Term | Chisq | Df | Pr(>F) |
|---|---|---|---|
| MY | 118.5869 | 2 | < 2.2e-16 *** |
| Habitat | 0.2924 | 1 | 0.5887 |

**Table S16: Emmeans Contrasts for Model 5.**

| | Odds ratio | SE | Df | z ratio | P value |
|---|---|---|---|---|---|
| Summer 2023-Summer 2024 | 0.01366 | 0.00596 | 1 | -9.833 | <.0001 |
| Summer 2023-Winter 2024 | 0.00871 | 0.00380 | 1 | -10.884 | <.0001 |
| Summer 2024-Winter 2024 | 0.63788 | 0.11400 | 1 | -2.515 | 0.0319 |

Results averaged over the levels of Scenario, Degrees of freedom method : Kenward-roger. Results are given on the log scale (not the response one).

**Fig S15: Diagnostic plots for Model 5.**

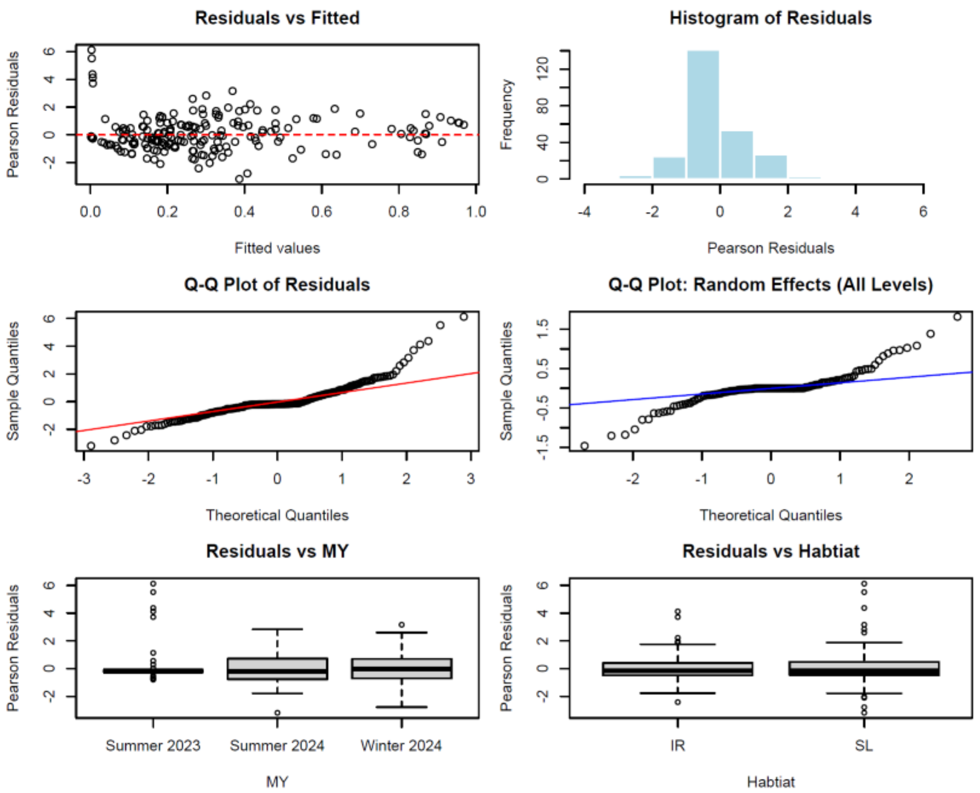

### Details for Model 6.1

**Table S17: Analysis of Deviance, Type II Wald F Tests for Model 6.1**

| Term | F | Df | Pr(>F) |
|---|---|---|---|
| Year | 86.1539 | 1 | < 2.2e-16 *** |
| Season | 7.9278 | 1 | 0.005185 ** |
| Year:Season | 0.0824 | 1 | 0.774209 |

**Fig S16: Fig. Diagnostic plots for Model 6.1**

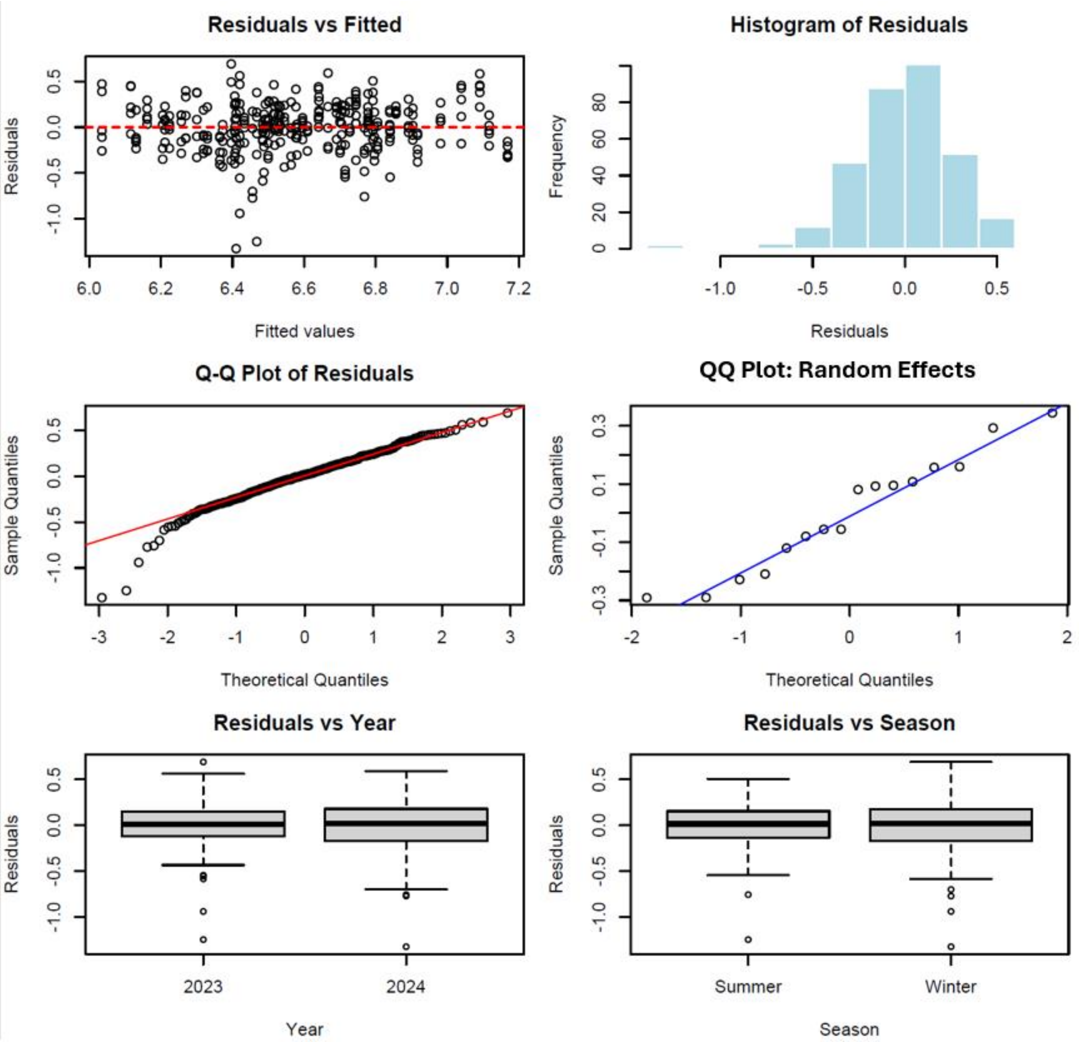

### Details for Model 6.2

**Table S18: Analysis of Deviance, Type II Wald F Tests for Model 6.2.**

| Term | F | Df | Pr(>F) |
|---|---|---|---|
| MY | 20.752 | 3 | 7.953e-13 *** |
| Type | 536.920 | 1 | < 2.2e-16 *** |
| MY:Type | 12.719 | 3 | 4.459e-08 *** |

**Table S19: Emmeans Contrasts for model 6.2.**

| Contrast | Estimate | SE | t.ratio | p-value |
|---|---|---|---|---|
| **Type Resident** | | | | |
| Summer 2023 – Summer 2024 | -0.39461 | 0.0708 | -5.577 | <.0001 |
| Summer 2023 – Winter 2023 | 0.24255 | 0.0693 | 3.498 | 0.0028 |
| Summer 2023 – Winter 2024 | -0.25050 | 0.0703 | -3.563 | 0.0022 |
| Summer 2024 – Winter 2023 | 0.63716 | 0.0700 | 9.101 | <.0001 |
| Summer 2024 – Winter 2024 | 0.14411 | 0.0710 | 2.031 | 0.1778 |
| Winter 2023 – Winter 2024 | -0.49304 | 0.0695 | -7.091 | <.0001 |
| **Type Visitor** | | | | |
| Summer 2023 – Summer 2024 | -0.11380 | 0.0710 | -1.604 | 0.3773 |
| Summer 2023 – Winter 2023 | -0.05626 | 0.0695 | -0.809 | 0.8503 |
| Summer 2023 – Winter 2024 | 0.10741 | 0.0707 | -1.518 | 0.4271 |
| Summer 2024 – Winter 2023 | 0.05754 | 0.0700 | 0.822 | 0.8441 |
| Summer 2024 – Winter 2024 | 0.00639 | 0.0712 | 0.090 | 0.9997 |
| Winter 2023 – Winter 2024 | -0.05115 | 0.0697 | -0.733 | 0.8336 |

**Fig S17: Diagnostic plots for Model 6.2.**

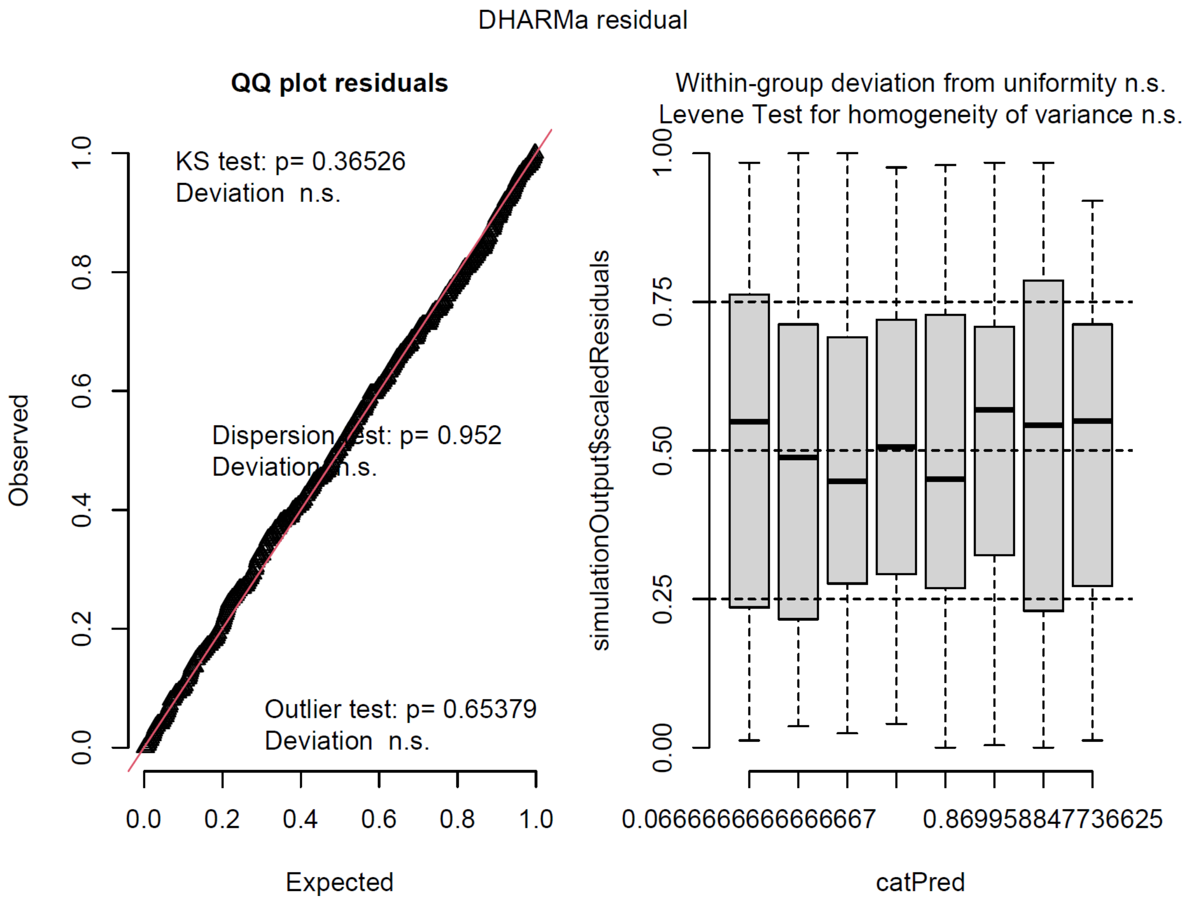

### Details for Model 7

**Table S20: Analysis of Deviance, Chi-square Tests for Model 7.**

| Term | Chisq | Df | Pr(>F) |
|---|---|---|---|
| Size | 0.0002 | 1 | 0.9894 |
| Sex | 26.7900 | 1 | 2.268e-07*** |
| Size:Sex | 0.0727 | 1 | 0.7860 |

**Fig S18: Diagnostic plots for Model 7.**

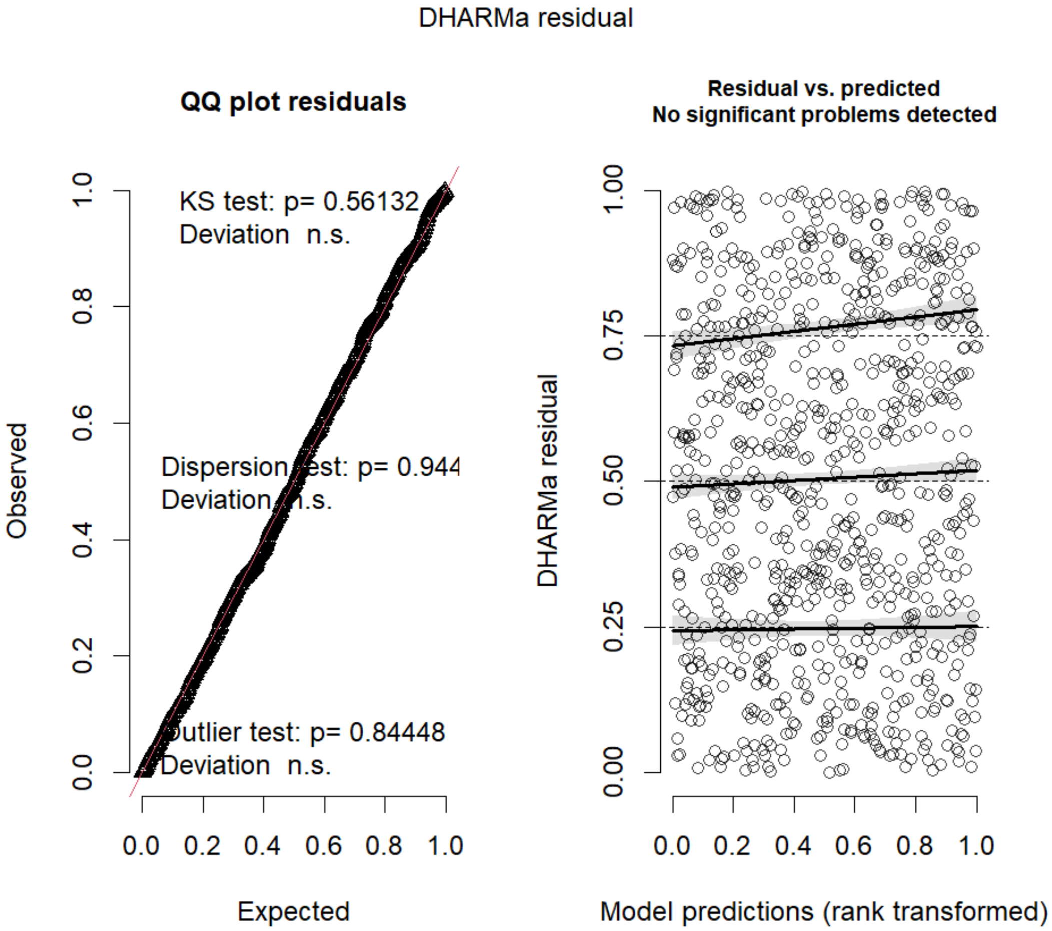
